# Host genetic background regulates the capacity for anti-tumor antibody-dependent phagocytosis

**DOI:** 10.1101/2023.05.09.540046

**Authors:** James E. Glassbrook, Justin B. Hackett, Maria C. Muñiz, Madeline Bross, Greg Dyson, Nasrin Movahhedin, Alexander Ullrich, Heather M. Gibson

**Affiliations:** Department of Oncology, Wayne State University School of Medicine, Karmanos Cancer Institute, Detroit, MI; Department of Biochemistry Microbiology and Immunology, Wayne State University, Detroit, MI

**Keywords:** Diversity Outbred, Anti-tumor Antibody, Targeted Immunotherapy, Antibody-Dependent Cellular Phagocytosis, scRNA-seq, Collaborative Cross

## Abstract

**Background:** Antitumor antibody, or targeted immunotherapy, has revolutionized cancer treatment and markedly improved patient outcomes. A prime example is the monoclonal antibody (mAb) trastuzumab, which targets human epidermal growth factor receptor 2 (HER2). However, like many targeted immunotherapies, only a subset of patients benefit from trastuzumab long-term. In addition to tumor-intrinsic factors, we hypothesize that host genetics may influence subsequent immune activation.

**Methods:** To model the human population, we produced F1 crosses of genetically heterogeneous Diversity Outbred (DO) mice with BALB/c mice (DOCF1). Distinct DOCF1 mice were orthotopically implanted with the BALB/c-syngeneic TUBO mammary tumor line, which expresses the HER2 ortholog rat neu. Treatment with anti-neu mAb clone 7.16.4 began once tumors reached ∼200 mm^3^. Genetic linkage and quantitative trait locus (QTL) effects analyses in R/qtl2 identified loci associated with tumor growth rates. Locus validation was performed with BALB/c F1 crosses with recombinant-inbred Collaborative Cross (CC) strains selected for therapy-associated driver genetics (CCxCF1). The respective roles of natural killer (NK) cells and macrophages were investigated by selective depletion in vivo. Ex vivo macrophage antibody-dependent phagocytosis (ADCP) assays were evaluated by confocal microscopy using 7.16.4-opsonized E2Crimson-expressing TUBO tumor cells.

**Results:** We observed a divergent response to anti-tumor antibody therapy in DOCF1 mice. Genetic linkage analysis detected a locus on chromosome 10 that correlates to a robust response to therapy, which was validated in CCxCF1 models. Single-cell RNA sequencing of tumors from responder and non-responder models identified key differences in tumor immune infiltrate composition, particularly within macrophage (Mφ) subsets. This is further supported by *ex vivo* analysis showing Mφ ADCP capacity correlates to *in vivo* treatment outcomes in both DOCF1 and CCxCF1 models.

**Conclusions:** Host genetics play a key regulatory role in targeted immunotherapy outcomes, and putative causal genes are identified in murine chromosome 10 which may govern Mφ function during ADCP.

## 1 Background

Monoclonal antibody (mAb) therapies have drastically improved cancer patient outcomes over the last 20 years. Trastuzumab is often lauded as one of the great success stories in mAb therapy as the standard of care for HER2^+^ metastatic and early-stage breast cancers as well as other HER2^+^ malignancies [1]. As adjuvant therapy in combination with chemotherapeutics, response rates are >75% [2]. Despite this success many patients that initially respond to trastuzumab monotherapy progress, suggesting that intrinsic and acquired resistance are common [3]. HER2 is a tyrosine kinase in the epidermal growth factor receptor (EGFR) family involved in cellular proliferation and survival. It is highly expressed in a variety of cancers, including 20-30% of breast carcinomas [4]. Successful induction of endogenous immunity is key for an effective response to targeted mAb therapy, but it is unclear why some patients are able to mount this response while others fail. The association of increased levels of stromal and tumor infiltrating lymphocytes (TIL) to the tumor microenvironment (TME) with an increase in survival has been reported [5,6]. Here we investigate the genetic basis of clinical resistance to targeted immunotherapy, focusing on regulation of immune-mediated killing. Therapeutic mAb effector functions are largely driven by innate immune cells including natural killer (NK) cells and macrophage (Mφ) which can kill tumor via antibody-dependent cellular cytotoxicity (ADCC) and antibody-dependent cellular phagocytosis (ADCP), respectively [7,8].

Early experiments implicated the mechanism of action of trastuzumab to be ADCC, although these were performed in purified human peripheral blood mononuclear cells (PBMC), and no distinction was made between ADCC and ADCP [9]. Elevated levels of NK cells are seen in trastuzumab-treated HER2^+^ breast cancer in humans; and while levels of Mφ are not statistically different between treated and control groups, infiltration is increased in the peripheral margins of residual tumor [10]. Both ADCC and ADCP are mediated by the recognition of antibody via members of the F_c_γR family. The binding properties of the F_c_γR family differ significantly between human and murine systems; however all F_c_γR in mice recognize IgG2a, similarly to human receptors recognizing subclass IgG1.

ADCC is governed by the recognition of the F_c_ region of antibody (Ab) on opsonized target cells via F_c_γRIII (CD16) on NK cells in both mice and humans, culminating in the release of perforin and granzyme-filled granules which mediate cell killing [11]. CD16, however, is also expressed by a subset of human and murine monocytes that have also been shown to be capable of the process [12]. ADCP is thought to mainly be carried out by Mφ, although any phagocytic cell is capable. This process is well described [13]. Murine receptors include high affinity pro-phagocytosis F_c_γRI (CD64) and F_c_γRIV (CD16-2), pro-phagocytosis low-affinity F_c_γRIII (CD16), and anti-phagocytosis F_c_γRIIb (CD32b) [14]. Engagement of F_c_γR with an opsonized target initiates a signaling cascade that triggers the stepwise process of ADCP, and a chain of events core to our understanding of how the innate immune system interfaces with the adaptive arm. Adaptive immunity may be required for complete tumor regression after targeted mAb treatment [15] and CD8^+^ T_c_ cell depletion is detrimental to survival in multiple models [16,17].

The genetic regulators of these processes are difficult to study by traditional means. To address the dearth of pre-clinical genetic models of genetic diversity, the Diversity Outbred (DO) population was developed. This was accomplished by the successive outbreeding of eight inbred strains, resulting in a population of genetically unique mice [18]. A subset were re-inbred to create dozens of Collaborative Cross (CC) lines. This model coupled with two genetic tools (GigaMUGA and R/qtl2) allows us to couple driver loci with a given phenotype of interest [19]. Some germline mutations have been reported to associate with HER2^+^ breast cancer susceptibility to trastuzumab [20] however we are using a targeted therapy that does not directly induce apoptosis. As a result, this model may more broadly apply to targeted immunotherapeutics generally. Given the lingering problem of innate resistance to targeted mAb therapies, we set out to pinpoint the host genetic basis of mounting a successful response to therapy. We identify a locus on chromosome (Chr)10 that correlates to response, confirm that locus in clonal models, and identify ADCP as a key innate immune process influenced by this locus that may have the potential for clinical intervention.

## 2 Materials and methods

### 2.1 Mice

Male and female mice were utilized in ∼1:1 ratio. Age- and sex-matched BALB/c mice and DO dams (6-8 weeks old) were purchased from The Jackson Laboratory (Bar Harbor, ME). Naming was carried out in accordance with the International Committee on Standardized Genetic Nomenclature for Mice. CC mice were purchased from the Systems Genetics Core Facility at University of North Carolina (UNC). CC lines used include the following: CC006/TauUnc, CC010/GeniUnc, CC024/GeniUnc, CC031/GeniUnc, and CC035/Unc [21–24]. Two to four DOxBALB/c F1 (DOCF1) siblings per dam were included in treated cohorts. One sibling from each litter was incorporated into the untreated cohort. Colony maintenance was performed under the supervision of Wayne State University (WSU) Division of Laboratory Animal Resources (DLAR). All animal procedures were approved by the WSU Institutional Animal Care and Use Committee. Tumor growth experiments were performed on mice 6–8 weeks old with inoculations administered in the 4^th^ inguinal mammary fat pad using ∼2×10^5^ TUBO cells suspended in DMEM (Gibco, 11965092). Tumors were measured 3 times/week (V = (W×W×L)/2).

### 2.2 Cell Lines and mAb Targets

TUBO cells were provided by Professor Wei-Zen Wei at WSU. B16F0 cells were acquired from American Type Culture Collection (ATCC, CRL-6322). TUBO/Crimson and B16/Crimson cell lines were generated in house via lipofectamine (Invitrogen, 11668-027) transfection of plasmid (Addgene, pEF.myc.ER-E2Crimson). Verification of E2Crimson expression of TUBO/Crimson and B16/Crimson was performed via flow cytometry (FC) with the help of WSU’s Microscopy, Imaging & Cytometry Resources (MICR). NCTC clone 929 (L929, Sigma, 85011425) were used to generate conditioned medium with M-CSF (Sigma Aldrich, 85011425). Cells were cultured at 37°C and 5% CO_2_ in a sterile environment. TUBO and TUBO/Crimson were cultured using high-glucose DMEM base medium with 10% heat-inactivated fetal bovine serum (FBS) (R&D Systems, S11150H), 10% NCTC-109 medium (Invitrogen, 21340039), 2 mM L-glutamine (Invitrogen, 25030081), 0.1 mM MEM (Invitrogen, 11140050), 1% penicillin (100 U/mL), 1% streptomycin (100 μg/mL) (P/S). E2Crimson expression was maintained via 600 μg/mL Geneticin G418 (Invivogen, ant-gn-1), and 7.5% sodium bicarbonate (NaHCO_3_) (Gibco, 25080-094). B16 and B16/Crimson cells were cultured in DMEM (10% FBS, 1% P/S). L929 cells were grown in RPMI (Gibco, 72400-047) supplemented with 10% FBS and P/S (R10).

Bone marrow (BM) cells were collected upon sacrifice and isolated as previously described [25]. BM cells were differentiated into bone marrow-derived Mφ (BMDM) in culture on non-tissue culture treated petri dishes using 30% L929 conditioned R10 media for a period of 7-9 days. Anti-neu mAb clone 7.16.4. was produced by Envigo in nude mice via hybridoma. Anti-TYRP1 mAb clone TA99 was acquired from Bio X Cell (BE0151). When used as opsonizing agents for confocal microscopy/FC, TUBO/B16 cells were incubated for 20 minutes at 4°C at a mAb concentration of 5µg/mL.

### 2.3 Cell Depletion

NK cell depletion was accomplished via anti-Asialo-GM1 (AsGM1) (Biolegend, Poly21460), as previously described [26]. Briefly, CCxCF1 mice received 2 doses of 50µL anti-AsGM1 intraperitoneally 2 days prior to mAb therapy and 1 week later. To deplete Mφ, 400µg mAb anti-CSF1R (Bio X Cell, BE0213) was given two days prior to treatment, and every two days thereafter as described previously [27].

### 2.4 Immunohistochemistry (IHC)

IHC was performed using primary antibodies CD86 (1:800, tris, Invitrogen, PA5-114995), Arg1 (1:4000, citrate, Proteintech, 16001-1-AP), and CD8α (1:400, tris, Cell Signaling Technology, 98941). Tumors were harvested, formalin fixed and paraffin embedded (FFPE), and sectioned at 5 µm. After deparaffinization in xylene and graded ethanol, antigen retrieval was performed using tris buffer at pH 9 (CD86, CD8α) or citrate at pH 4 (Arg1). Blocking was performed using BLOXALL (Vector Laboratories, SP-6000). Primary antibody incubation was performed overnight at 4° C in a humidified chamber. Secondary antibody Horse anti-Rabbit IgG, ImmPRESS Alkaline Phosphatase (AP) (Vector Laboratories, MP-5401) was performed for 30 minutes. Detection was performed via Vector Red AP Kit (Vector Laboratories, SK-5100). Enumeration was conducted using Trainable Weka Segmentation, available through the FIJI (FIJI is just ImageJ) [28–30]. Quantification was performed on three representative 40x fields per tumor.

### 2.5 Phagocytosis Assays and Confocal Microscopy

ADCP assays were carried out using BMDM in chamber slides (Corning, 354114) in co-culture with mAb clone 7.16.4-opsonized TUBO/Crimson or TA99-opsonized B16/Crimson (20k BMDM:20k tumor cells) versus non-opsonized controls. After 4 hours of co-culture cells were fixed with 4% paraformaldehyde (Sigma-Aldrich, 158127) and permeabilized with 0.1% Triton X-100 Surfact-Amps Detergent (Thermo Scientific, 85111). Staining was performed for: EEA1 (Abcam, ab206860), 5µg/mL overnight at 4° C and secondary (Abcam, ab150129) at 2µg/mL, 45 minutes at room temperature (RT); F4/80 (Invitrogen, PA5-32399), 1:100 overnight at 4° C, and secondary (Abcam, ab150064) at 2µg/mL, 1 hour at RT. Nuclear staining/mounting was performed using ProLong Gold Antifade Mountant (Invitrogen, P36931). Images were taken on a Zeiss-780 confocal microscope, with the support of WSU’s MICR Core. Three 40x fields were taken for each experimental well, and one for each control. Image analysis was performed via FIJI. One Mφ with high uptake was chosen from each field, outlined using EEA1 staining as reference for cell area, and E2Crimson uptake was reported as MFI minus background. The antibody-independent Vybrant Phagocytosis Assay Kit was used per manufacturer instruction (Invitrogen, V6694).

### 2.6 Flow Cytometry

FC assays were carried out using BD LSR II and Northern Lights flow cytometers with the assistance of WSU’s MICR Core. Zombie near infrared (NIR) fixable viability stain (BioLegend, 423105) was used at a dilution of 1:1000 (15 minutes). Antibodies include: CD64 (Invitrogen, 12-0641-82), CD32b (Invitrogen, 17-0321-82), CD16 (Invitrogen, MA5-40921), and CD16-2 (Invitrogen, MA5-28253). Staining was performed at 1:200 for 30 minutes prior to fixation overnight in 1% phosphate buffered formalin (Fisher Scientific, SF100-4). Note: CD64 mAb clone (X54-5/7.1) will not detect allelic variant d (present in the NOD background), no DOCF1 mice included in FC experiments were NOD at the CD64 locus.

### 2.7 qRT-PCR

Tumor tissues were flash frozen in liquid nitrogen upon collection. RNA was extracted via TRIzol (Thermo Fisher, 15596018) post homogenization by Tissue Tearor (Biospec, 985370-04). cDNA was prepared using ProtoScript II reverse transcriptase (New England Biolabs, M0368S). Quantitative reverse-transcribed PCR (qRT-PCR) was performed using iTaq Universal Probes Supermix (Bio-Rad Laboratories, 1725131), 5 ng cDNA/well and 500 nM primers (**Supplemental Table 1**). Quantification was reported as difference in cycle threshold (ΔCT) relative to GAPDH x 1×10^6^.

### 2.8 DO Genotyping, Linkage and Quantitative Trait Locus Analysis

DOCF1 mouse genotyping was performed via GigaMUGA [31], by Neogen. Linkage and QTL analysis were performed in R/qtl2 (v0.24) in R version 4.0.3. DOCF1 haplotypes were reconstructed using a hidden Markov model. Linkage analysis employed a mixed effects linear model with sex and relatedness (determined via kinship matrix) as co-factors. Log of odds (LOD) scores >6 were considered for downstream analysis, and single strain Quantitative Trait Locus (QTL) effects were examined to determine putative positive/negative drivers. Scripts and R session data available upon request.

CC strains were selected by strain background ± 1 megabase pairs (Mbp) of the Chr10 locus. Alternative prospective peaks were cross-referenced to ensure CC strains did not contain positive/negative driver backgrounds at other putative loci (Chr 3 and 11). CC mouse genomes, status, and ideograms provided and maintained by UNC Computational Systems Biology.

### 2.9 Quant-seq

Snap frozen tumor collected from 8 high tumor doubling time (TDT) samples (DOCF1), 8 low-TDT samples (DOCF1), and 2 CC024CF1 samples were utilized for bulk transcriptomics. Quant-seq Library Preparation was performed by WSU’s Genome Sciences Core Facility using the QuantSeq 3’ mRNA-Seq Library Prep Kit FWD. High Sensitivity D1000 ScreenTape peak ranges between 100 bp and 700 bp in accordance with Lexogen guidelines. BBduk parameters available upon request. Counts were generated for each gene region and differential gene expression analysis was used to compare transcriptome changes between conditions. Significantly altered genes (|log fold change| ≥ 2; p-value ≤ 0.05) were used for downstream pathways analysis. Gene ontology (GO) analysis was performed using DAVID (https://david.ncifcrf.gov/) [32] and Cytoscape v3.7.2 [33], primarily using the Biological Process ontology.

### 2.10 scRNA-seq

Tumors were dissociated (Miltenyi, 130-096-730) and CD45^+^ enriched (StemCell Technologies, 100-0350). Replicates were tagged using the 10x Genomics 3’ CellPlex Cell Multiplexing Oligo (CMO) system (10x Genomics, PN-1000261) with library preparation using the Chromium Next GEM Single Cell 3’ Kit (10x Genomics, PN-1000268) and sequencing was performed on the NovaSeq 6000 (Illumina) by the WSU GSC. Run parameters and quality control metrics recommended by Illumina and 10x Genomics were all met or exceeded. Alignment was performed using the Cellranger pipeline (GSC Alignment refdata-cellranger-mm10-3.0.0).

Downstream data analyses were performed using Seurat in R v4.2.0 [34]. Cell type identification was performed via SingleR [35], with adjustments based on canonical marker genes. Inclusion criteria were defined as follows (>200 and <2500 genes expressed, <10k RNA counts, <5% mitochondrial genes content). Over 60k cells were included in downstream analyses. Cells were clustered on the top 2,000 most variable genes. Doublet exclusion was achieved by removing cells with >1 CMO tag, as well as manual exclusion of clusters with abnormally high read counts *post hoc*.

### 2.11 Other Data and Statistical Analysis

Statistical analyses were conducted using Graphpad Prism v. 9.3.1, including growth curves, survival analyses, normality and distribution statistics, χ^2^ testing for proportions, rank correlations, analysis of variance (ANOVA) of means, and several t-test variations. Data are presented as the mean ± S.D. unless otherwise noted. A p-value < 0.05 is considered statistically significant.

## 3 Results

### 3.1 Host genetic background as a predictor of response to targeted mAb therapy

To model the influence of host genetics upon response to targeted immunotherapy, we produced a cohort of DOCF1 mice (n = 126) from 35 distinct DO dams (**Figure 1A**). DOCF1 mice were orthotopically implanted with rat neu^+^ (a HER2 ortholog) TUBO mammary carcinoma cells [36]. As our model targeted immunotherapy, we selected mAb clone 7.16.4, which binds to the extracellular domain of neu with minimal cytotoxic effects *in vitro* [16]. mAb 7.16.4 is a mouse IgG2a isotype, with similar immune effector engagement as a human IgG1, such as trastuzumab. Upon reaching a volume of ∼150 mm^3^, treatment with 7.16.4 therapy was initiated (**Figure 1B**). At day 34, ∼45% of mice had completely responded (CR) to therapy, eliminating primary tumor. These mice also rejected a contralateral re-challenge on day 45 (without additional therapeutic intervention), suggesting protective adaptive anti-tumor immune induction (**Figure 1C**). The remaining ∼55% of the cohort partially responded (PR) to therapy to varying degrees (**Figure 1D**). Untreated control mice reliably accepted tumor and succumbed to disease at a similar rate, however it is important to note that ∼4% of DOCF1 mice rejected TUBO without therapeutic intervention and were excluded from the study (**Figure 1E**). To quantify efficacy, we performed regression modeling on each growth curve to determine TDT (18 to 94 days) (**Figure 1F**). TDT of CR mice was set to 200 days to distinguish from PR mice for linkage analysis. The proportion of CR cage-mates follows a gaussian distribution (**Supplement 1A**); however, we find that siblings respond similarly. Both 0% and 100% CR rates are inflated among littermates, suggesting variation in response is heritable and therefore genetic (**Supplement 1B**).

**Fig 1.**
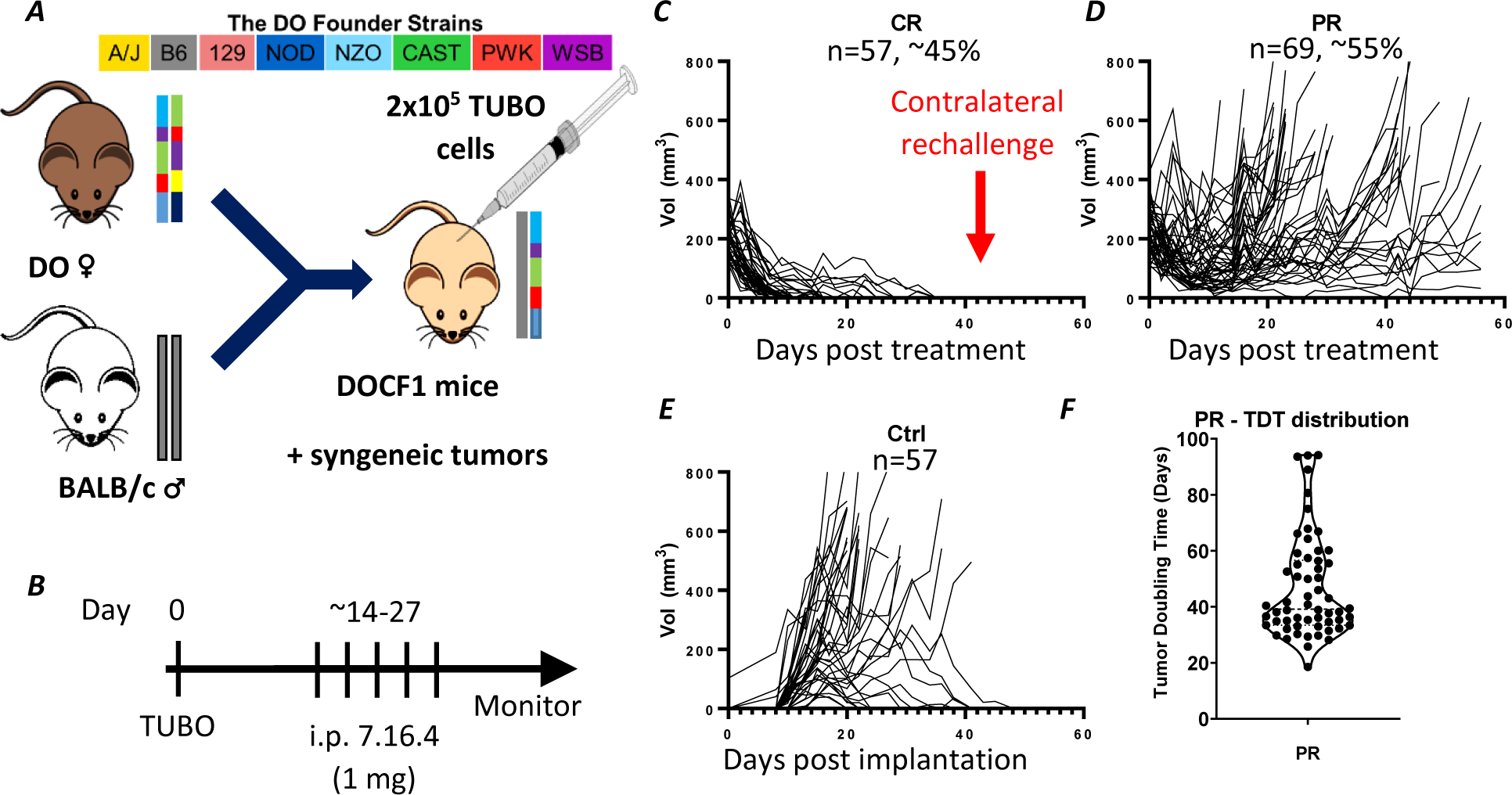
Diverse genetics yields wide variation in response to targeted immunotherapy. (***A***) Color scheme representing the 8 founder genomes comprising the DO mouse model, and breeding scheme for development of DOCF1 mice. (***B***) Treatment timecourse depicting TUBO tumor cell inoculation and administration of 5 doses of anti-neu mAb clone 7.16.4 (1 mg/dose) upon tumors reaching ∼150 mm^3^. Tumor growth from treatment initiation of (***C***) CR, wherein contralateral rechallenge of TUBO was implanted on day 45 (red arrow), and (***D***) PR. (***E***) Tumor growth curves of control (Ctrl) untreated mice from the time of inoculation. (***F***) Distribution of PR TDT.

All mice were genotyped using the GigaMUGA array [31], and genetic linkage analysis was performed with the R/qtl2 package [19] to evaluate TDT association against host genetics (**Figure 2A**). The most highly associated loci reside on Chr1 and 10; each with logarithm of odds (LOD) scores >6. The Chr1 locus yielded indecipherable QTL effects (not shown); however, the Chr10 locus (∼40.6-46.6 Mbp) clearly implicates PWK genetics as a positive phenotypic driver of a robust response to targeted therapy, and conversely A/J as a negative driver (**Figure 2B**). Among the DOCF1 cohort, all mice with PWK at Chr10 in this locus (n = 6) are among the CR mice. Loci on Chr 3 and 11 were also identified as potential regions of interest (**Supplement 2**).

**Fig 2.**
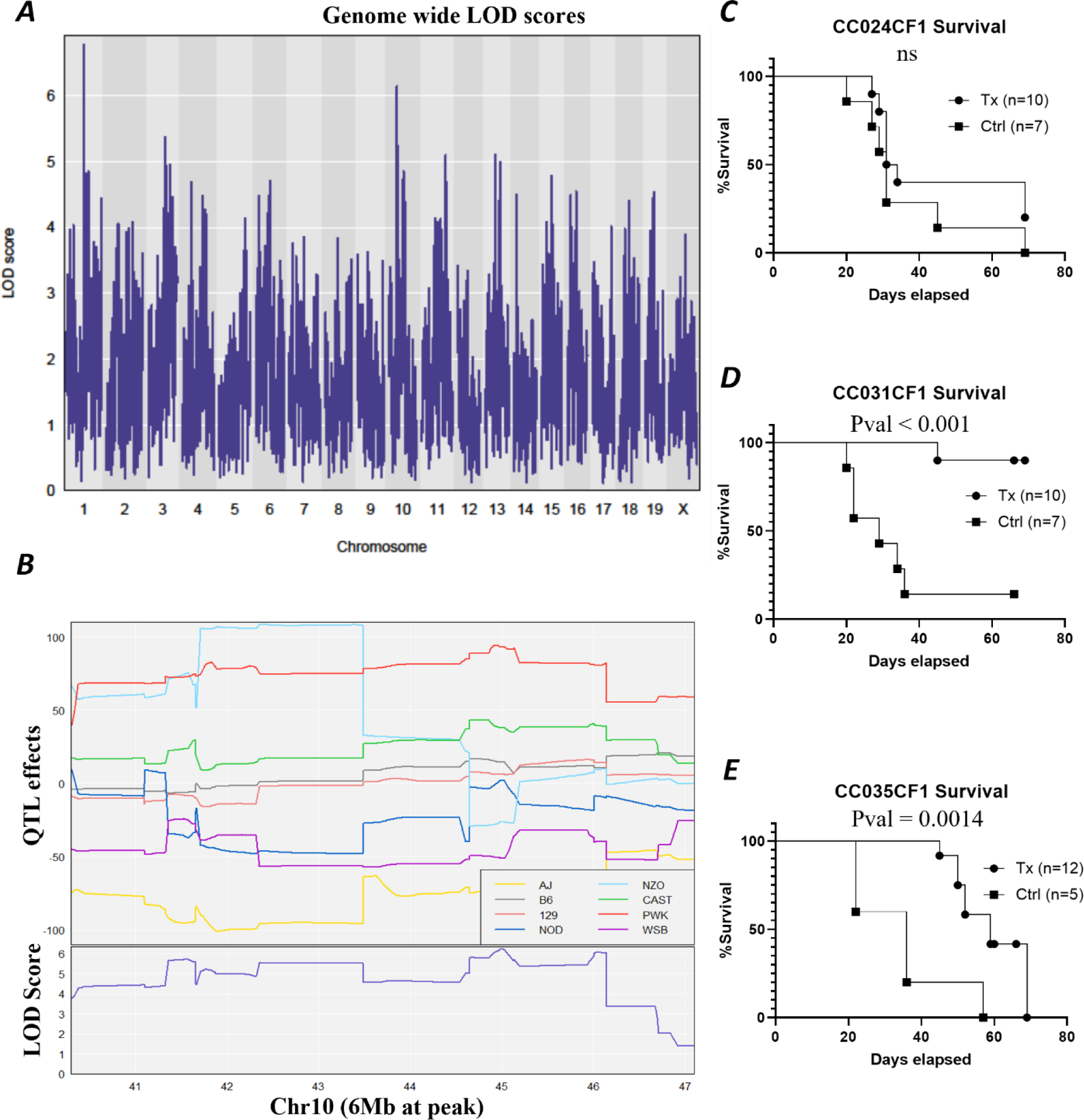
Genetic linkage analysis of targeted immunotherapy response in DOCF1 mice identifies loci associated with targeted immunotherapy response. ***(A)*** Genetic linkage analysis of DOCF1 mice in Fig 1 via R/qtl2 using TDT as an indicator of response to treatment. (***B***) Above: Quantitative Trait Locus (QTL) effects plot at the peak of interest in Chr10. Below: corresponding LOD scores. (***C***) Survival of predicted NR stain CC024CF1. (***D***) Survival of predicted CR stain CC031CF1. (***E***) Survival of predicted PR stain CC035CF1.

To validate the Chr10 locus as a therapeutic regulator, CC strains were identified based on driver genetics at this locus. To avoid confounding genetic factors, strains with drivers of the Chr 3 and 11 loci were omitted. CC031 and CC035 were selected as putative responders, and CC024 was selected as a putative non-responder. CC mice were crossed with BALB/c to produce cohorts of CCxCF1 mice (**Supplement 3A**). CC024 is mostly A/J at this locus on Chr10, CC031 is 100% PWK, and CC035 is ∼50% PWK and ∼50% WSB (**Supplement 3B**).

As predicted, CC024CF1 did not respond to therapeutic intervention, with most tumors quickly surpassing the euthanasia threshold volume (500 mm^3^) regardless of treatment (**Figure 2C, Supplement 3C**). Most treated CC031CF1 mice reliably eliminated tumor (**Figure 2D, Supplement 3D**) and CC035CF1 showed a PR to therapy, though less dramatically than CC031CF1 mice (**Figure 2E, Supplement 3E**). This locus spans 26 protein-coding and 50 non-protein-coding genes. Many of the protein-coding genes are known regulators of immune signaling and maturation (**Supplement 4**), including 7 transcription factors (Bend3, Hace1, Prdm1, Scml4, Foxo3, Mtres1, and Nr2e1). A genome-wide breakdown of founder strain contributions for all CC mice utilized in this study is included (**Supplement 5**).

### 3.2 Canonical effector cell populations are confirmed to influence targeted mAb response

Both NK cells and Mφ can mediate the effector functions of targeted mAb therapy. To definitively test if these cell types are impacting therapeutic response in our CC models, we selectively depleted NK cells (using anti-Asialo-GM1) or Mφ (using anti-Csf1r) in CR model CC031CF1. Unsurprisingly, both NK and Mφ depletion are detrimental to overall survival in the context of 7.16.4 therapy, suggesting these innate cell populations each play an important role in tumor elimination (**Supplement 6**). This finding is in accordance with previous work using targeted immunotherapy anti-TYRP1 (clone TA99) in the B16 melanoma model [17].

To further evaluate innate and adaptive immune response in the TME of our original DOCF1 cohort, we performed IHC on FFPE tumors. An example of IHC enumeration is provided in **Supplement 7**. While no significant difference is seen in general Mφ marker CD68 or M1-associated marker CD86 between low and high-TDT tumors (**Figure 3A-B**), we do see elevated numbers of cells positive for the M2-associated marker Arg1 in slower-growing tumors (pval = 0.0159) (**Figure 3C**). This increase in Arg1 expression may be induced by a smoldering, insufficient immune response wherein tumor growth outperforms anti-tumor immunity. Supporting this notion, we also find elevated levels of CD8^+^ T_c_ staining in high-TDT tumors (pval = 0.0011) (**Figure 3D**), implicating adaptive immune induction as a key factor in controlling tumor burden. As mAb therapies are unlikely to directly activate CD8^+^ T cells, these results suggest immune priming may occur as a result of innate immune activity.

**Fig 3.**
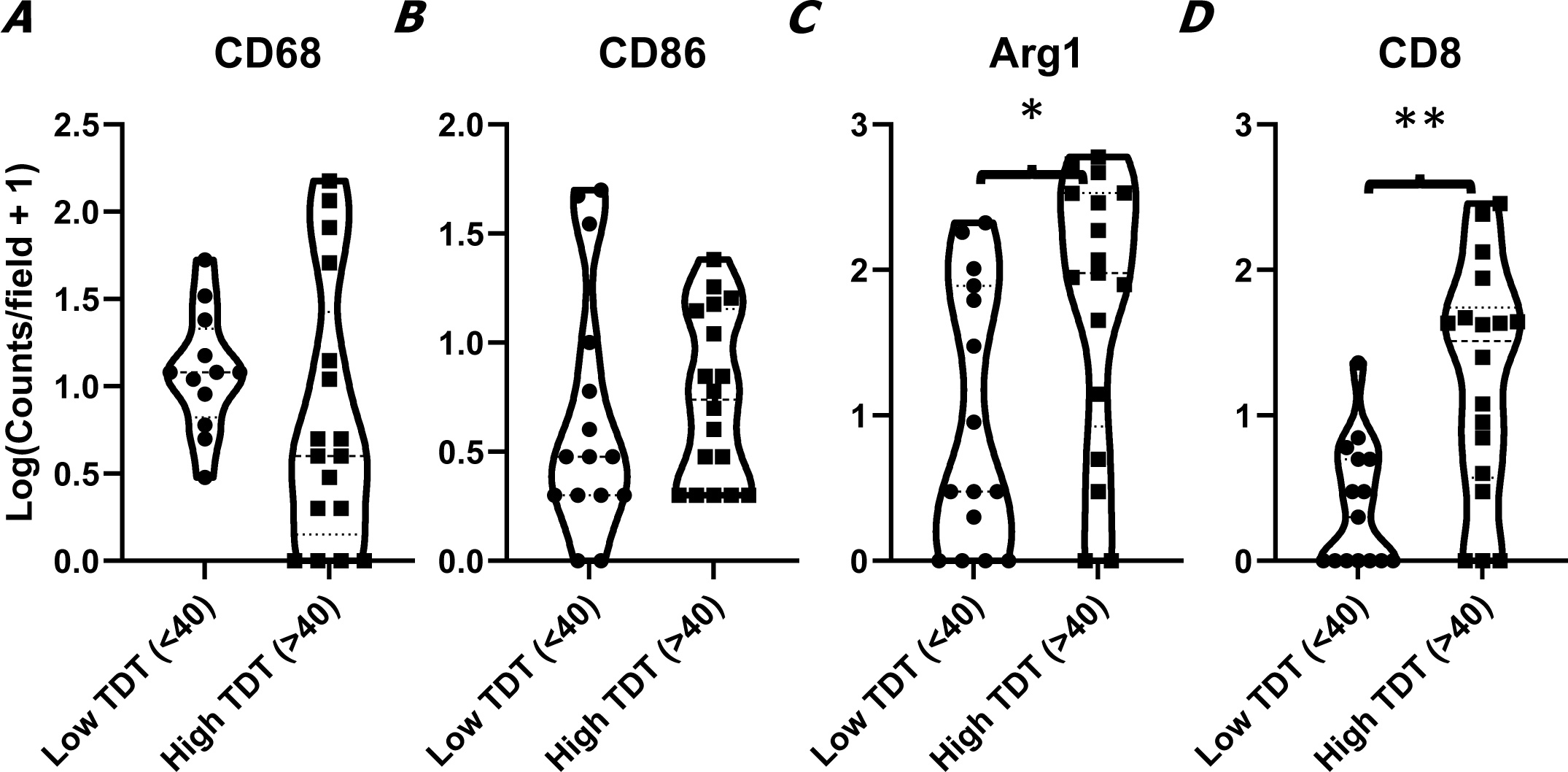
Relative abundance of key tumor infiltrating immune populations in the context of targeted immunotherapy. (***A-D***) Quantification of immune infiltrates in DOCF1 cohort. (***A***) CD68 (total Mφ). (***B***) CD86 (M1 Mφ). (***C***) Arg1 (M2 Mφ). (***D***) CD8 (CD8^+^ T_c_).

### 3.3 Transcriptomic analysis reveals transcriptional programs and immune infiltrate phenotypes that correlate with targeted immunotherapy response

To evaluate the transcriptome of the TME, we compared flash frozen tumor tissue from high-TDT (n=9) and low-TDT (n=9) DOCF1, and low-TDT CC024CF1 (n=2) mice, which were collected upon reaching the tumor threshold volume (>500 mm^3^). Similar to our IHC analyses, CR mice were unable to be evaluated due to a lack of remaining tumor tissue. Bulk mRNA-sequencing (Quant-seq) was employed, which identified differentially regulated transcripts based on the binary response: high (>40 days) vs low (≤40 days) TDT. A distance matrix was calculated based on each transcriptome’s relatedness (**Supplement 8**). In general, tumors with similar growth rates cluster together, however exceptions may suggest multiple transcriptional programs are in play that independently cause similar outcomes. This is also reflected in the top differentially regulated genes (**Figure 4A**). Interestingly, CC024CF1 samples cluster independently of the low-TDT DOCF1 samples, with unique transcriptional signatures. Gene set enrichment analysis implicates various biological processes; specifically, antigen processing/presentation, myeloid development/differentiation, and endocytosis/vesicular transport (**Figure 4B**). Many of these genes appear to be involved in innate immune signaling and ADCP, including endosomal and lysosomal membrane composition (**Supplement 9-10**).

**Fig 4.**
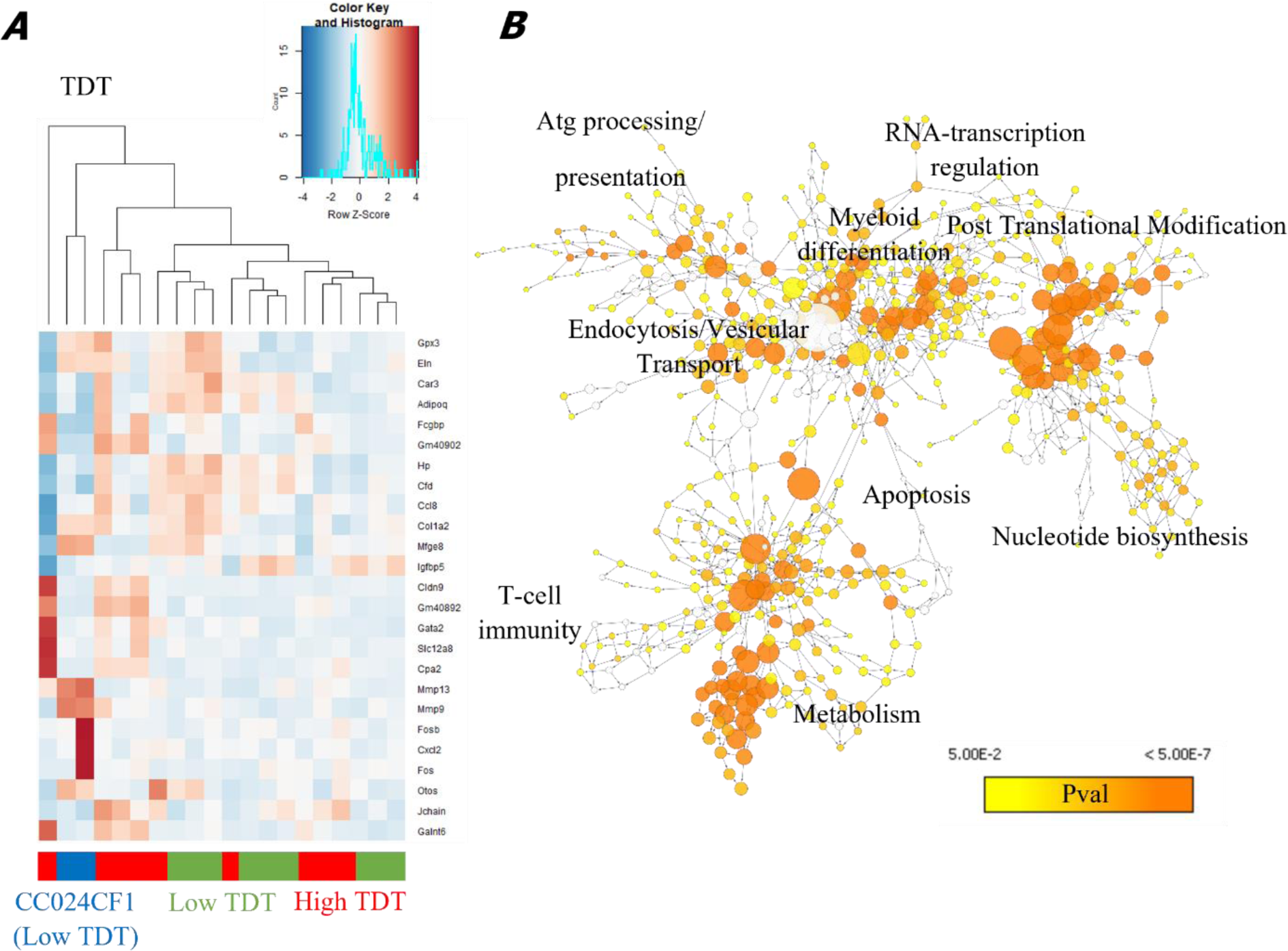
Transcriptomic profile of low-TDT vs high-TDT growing tumors reveals differentially expressed transcriptional programs. Bulk transcriptomic analysis of flash frozen TUBO tumor, implanted in DOCF1 and CC024CF1. *(**A**)* Top 25 differentially regulated genes, clustered by TDT. (***B)*** Over-represented GO terms (Biological Process) among differentially regulated genes.

Given the differential innate immune-related expression patterns in high and low-TDT tumors, we next sought to directly compare immune infiltrate phenotypes in tumors from clonal CCxCF1 models. Tumors from 7.16.4-treated (n=3 each) CC024CF1 (low-TDT), CC035CF1 (high-TDT), and CC031CF1 (CR) were harvested 29 days post tumor implantation, corresponding to 12 days after initiation of mAb therapy (after 3 doses). This partial course of treatment ensures sample collection at the therapeutic inflection point, before CR mice would otherwise eliminate tumor. scRNA-seq was performed on CD45^+^ enriched samples, capturing >60k cells.

Using data from the Immunological Genome Project [37] and R/SingleR [35], we compared the transcriptome of each cell to that of known purified cell populations. SingleR assigns a Spearman correlation score to each cell and each prospective cell type (**Supplement 11**) and designates an ID. Differences between each assigned ID from the median score of the group are depicted in a delta-plot and serve as a quality control metric (**Supplement 12**).

Following manual exclusion of doublets, we see the immune milieu is dominated by Mφ, T cells, NK cells, neutrophils, conventional dendritic cells (cDC), plasmacytoid dendritic cells (pDC), and B cells (**Figure 5A**). Heatmaps of the top 10 genes are found in **Supplement 13**. Cell identities largely align with predictions, although cDC, pDC, NK, T cell subsets, and actively mitotic cells were identified by canonical marker genes (**Supplement 13-14**). When we compare the relative proportions of immune cell populations between the treated low and high-TDT/CR strains, we see significant differences between groups by χ^2^ test for differences in proportions (with a Yates correction) and Monte-Carlo permutation test (**Figure 5B-C**). Neutrophils are overrepresented in low-TDT tumor infiltrate while monocytes, pDC, NK cells and B cells are overrepresented in high-TDT/CR strains.

**Fig 5.**
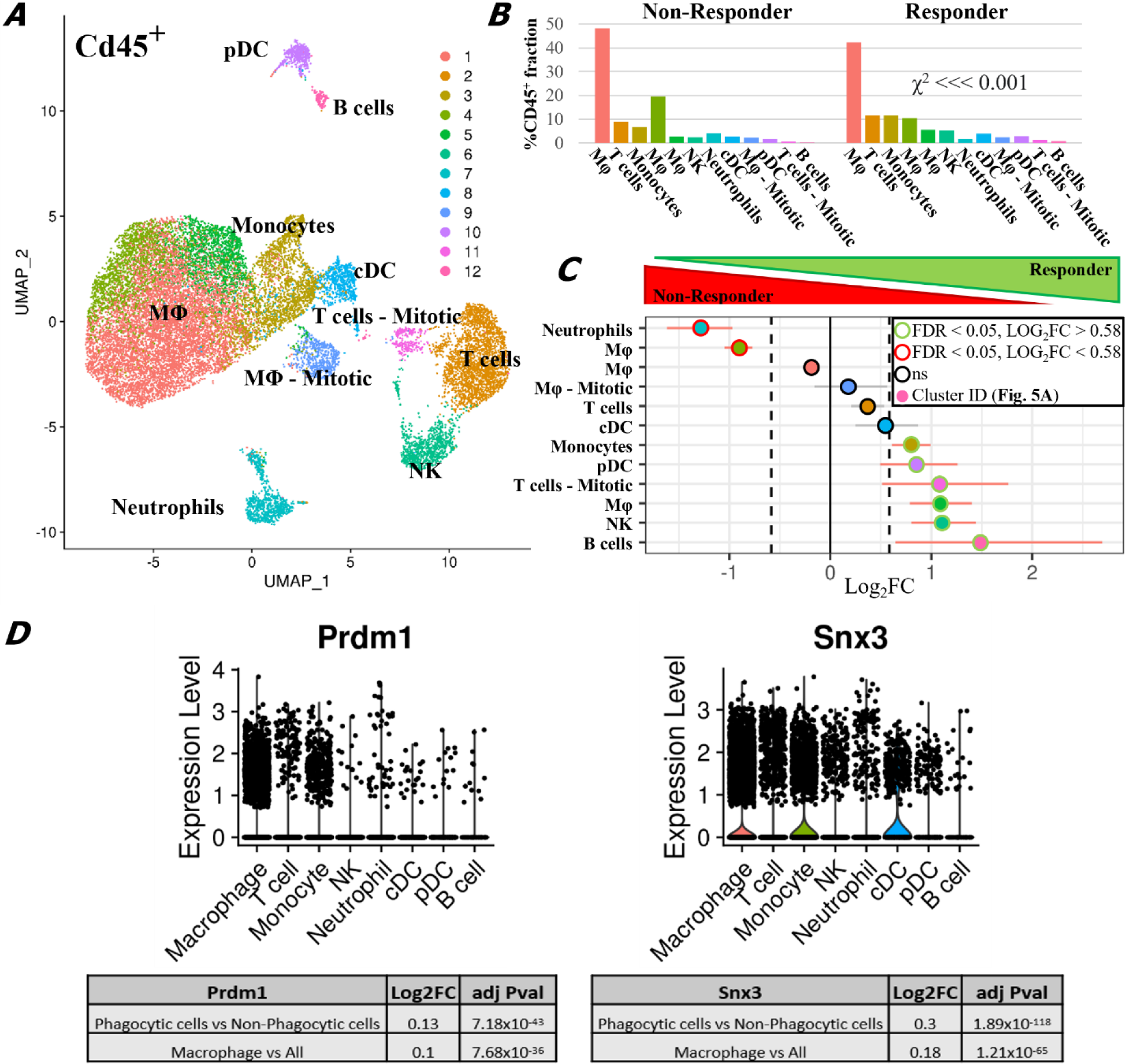
scRNA-seq reveals key differences in the immune milieu of low-TDT versus high-TDT/CR tumors after targeted immunotherapy. *(**A**)* UMAP dimensionality reduction, clustering CD45^+^ cells into 12 transcriptionally distinct clusters. *(**B**)* Relative abundance of CD45^+^ cells, comparing low-TDT and high-TDT/CR mice, χ^2^ test with Yates correction. ***(C)*** Fold change in abundance of identified clusters between low-TDT and high-TDT/CR mice. Significance testing by Monte Carlo permutation test. Log_2_ Fold change cut-off set to 0.58, ∼1.5 fold change. ***(D)*** Chr10 genes significantly expressed at higher levels in phagocytic cells vs non-phagocytic cells, and Mφ as a whole.

Because CC models were chosen based on their genetics at the Chr10 locus, we evaluated the expression patterns within the CD45^+^ fraction for all genes located within this locus. Only 19 of these genes are detectable in the scRNA-seq data, detailed by cell type in **Supplement 15**. We do not see any statistical differences in the transcription of these genes between low-TDT and high-TDT/CR models (not shown), suggesting SNPs in this region might not regulate the mRNA expression level, but instead could be involved in altered sequence, stability, or function of protein or transcript, or developmental processes prior to tumor infiltration. Importantly, genes from this locus tend to be more highly expressed in phagocytic cells, particularly Snx3 and Prdm1 in the Mφ compartment (**Figure 5D**). While NK cells are overrepresented in high-TDT/CR tumors, few transcriptional differences in these genes can be seen between conditions. Therefore, we sought to primarily investigate the phagocytic fraction.

Mφ contain more transcriptional heterogeneity than can be observed in the initial UMAP projection. Further subsetting of this population allows us to perform more detailed functional annotation, as well as assign more robust cell identities using canonical marker genes. When the phagocytic fraction is re-clustered, 4 subpopulations of Mφ (clusters 2, 3, 6 and 7) distinguish themselves from the core population (**Figure 6A**). Cluster 2 is enriched for genes involved in C1 complement secretion. Cluster 3 is characterized by genes known to promote proinflammatory processes and chemokines involved with neutrophil chemotaxis. Cluster 6 expresses genes known to regulate endocytosis, as well as several chemokines known to recruit monocytes such as CCL8. Both Clusters 3 and 6 were overrepresented in the high-TDT/CR strains, at more than double the frequency found in low-TDT tumors. Cluster 7, representing actively mitotic Mφ, is approximately equal between strains (**Figure 6A-C**). The top ten genes representing the transcriptional signatures of these subclusters are provided (**Supplement 16**).

**Fig 6.**
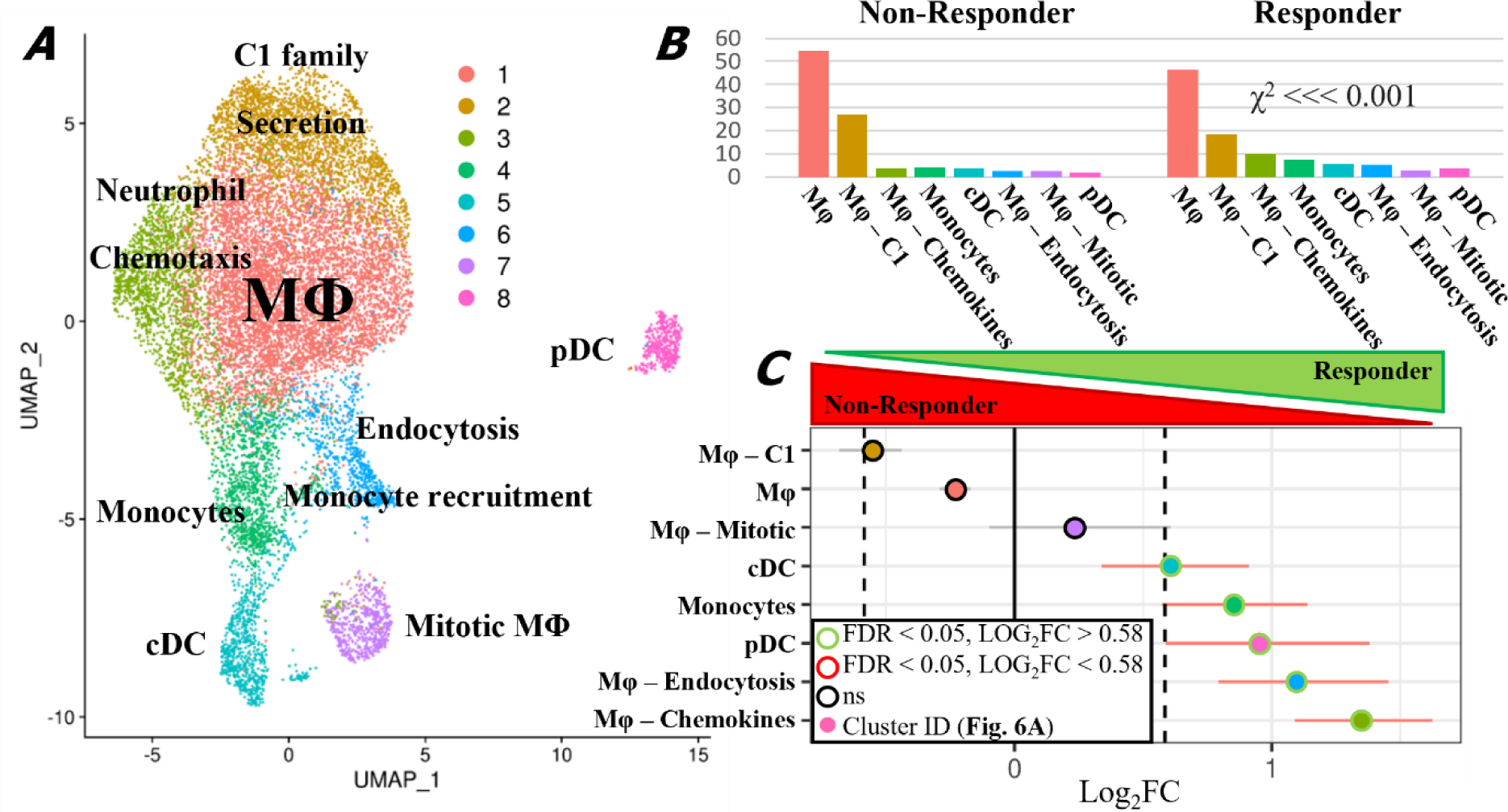
scRNA-seq subclustering of phagocytic fractions reveals key differential transcriptional profiles of Mφ between responder and non-responder models. *(A)* UMAP dimensionality reduction of phagocytic cells into eight transcriptionally distinct clusters. *(B)* Relative abundance of clusters, comparing low-TDT and high-TDT/CR mice. ***(C)*** Fold change in abundance of identified clusters between treated NR and PR/CR strains.

### 3.4 Capacity for ADCP predicates targeted immunotherapy outcomes

Given the transcriptional differences between the low and high-TDT/CR TME and the expression of Chr10 genes in the myeloid fraction, we proceeded with the hypothesis that ADCP may be regulated by the host genetic background at this locus. To investigate this further, BMDM were evaluated *ex vivo* from low-TDT (n=7), high-TDT (n=5), and CR (n=7) mice from the initial DOCF1 experiment. Mφ were cocultured with 7.16.4-opsonized TUBO cells stably transfected to express the pH-stable fluorescent protein E2Crimson (**Supplement 17A**), and tumor uptake within Mφ was evaluated by confocal microscopy. Samples lacking 7.16.4 served as a control for antibody-dependence. Monoculture BMDM were also stained for F4/80 to confirm successful Mφ differentiation (**Supplement 17B**). Quantification of E2Crimson was evaluated by outlining and subsequent measurement of E2Crimson MFI via FIJI (**Supplement 17C**).

BMDM generated from CR mice from the DOCF1 cohort exhibited the highest degree of E2Crimson uptake, showing many punctate vesicles containing E2Crimson. When we convert response level into ordinal numbers based on TDT and compare them via Spearman’s rank correlation, we find a significant trend toward higher uptake in BMDM collected from CR mice (pval = 8.994×10^−5^) with a ρ value = 0.66 (**Figure 7A**). The non-opsonized condition includes a mix of low and high-TDT mice, demonstrating the antibody-dependent nature of E2Crimson uptake. A representative image of a Mφ containing phagocytized E2Crimson can be seen in **Figure 7B**. This observation is recapitulated in the CCxCF1 mouse lines. CC models with the responder PWK genotype at the Chr10 locus more efficiently take up E2Crimson than mice with the A/J genotype (pval = 0.023) (**Figure 7C**). Importantly, this effect is not limited to cell line or mAb. When the same experiment is carried out using opsonized murine melanoma (B16F0) with mAb TA99 specific to antigen TYRP1, a similar trend is observed (pval = 0.0002) (**Supplement 18**). Representative images of Mφ derived from CC024CF1, CC031CF1, and CC035CF1 are shown in supplementary figures for both tumor lines TUBO/Crimson (**Supplement 19**) and B16/Crimson (**Supplement 20**).

**Fig 7.**
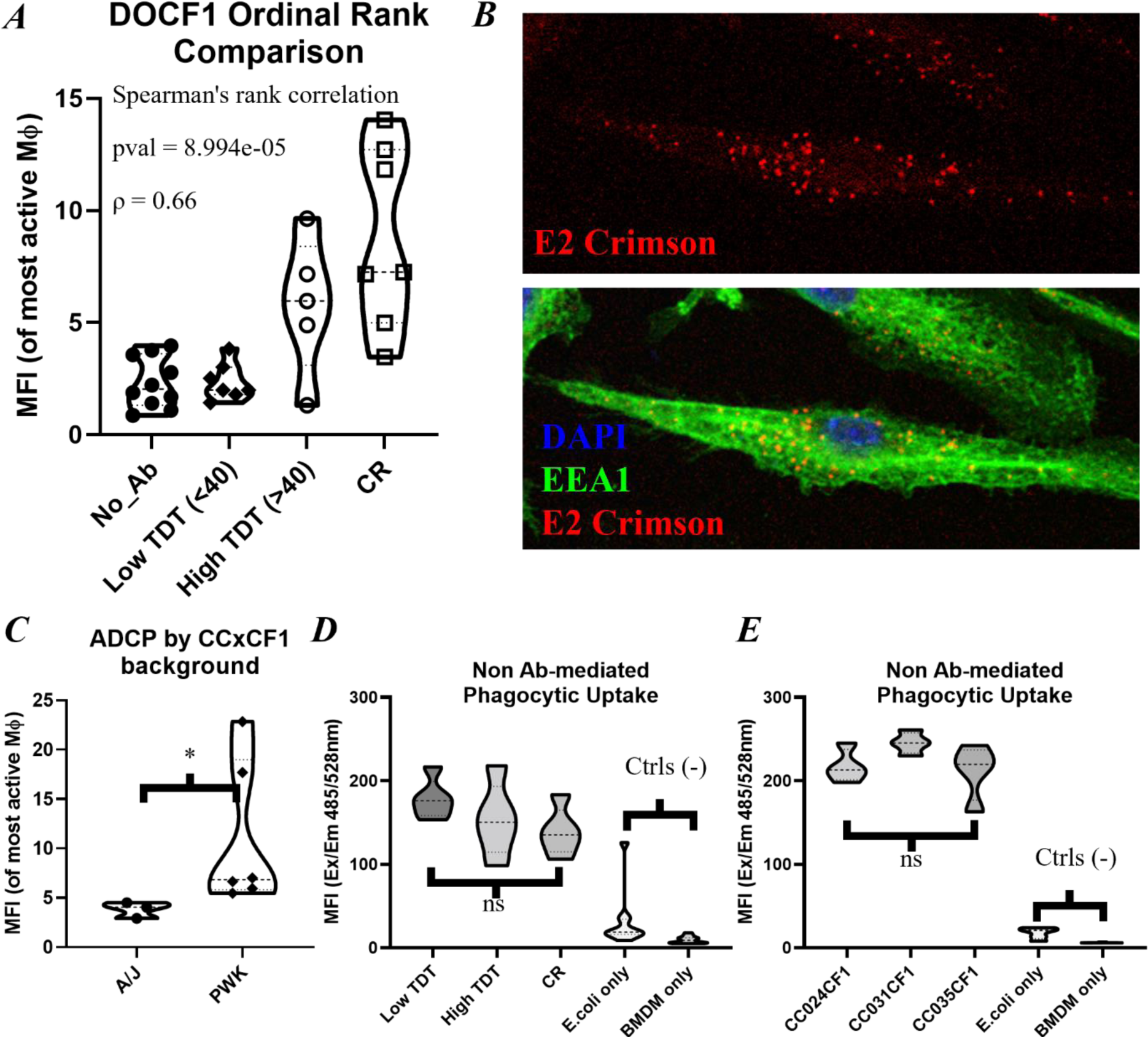
Host genetic background modulates BMDM capacity for ADCP. BMDM derived from DOCF1 and CCxCF1 in coculture with 7.16.4-opsonized TUBO/E2Crimson for 4 hours. *(**A**)* Ordinal rank comparison of representative E2Crimson uptake in BMDM derived from DOCF1 mice selected across three fields of view. *(**B**)* Representative image of BMDM differentiated from a CR mouse. DAPI (blue), EEA1 (Green), E2Crimson (Red). Above – E2Crimson only. Below – composite image. *(**C**)* E2Crimson MFI in representative BMDM derived from CCxCF1 mice across three fields of view, grouped by genetic background at Chr10 locus of interest. *(**D-E**)* E2Crimson MFI of BMDM in coculture with fluorescent E. coli. *(**D**)* DOCF1 *(**E**)* CC024CF1 (low-TDT), CC031CF1 (CR), and CC035CF1 (high-TDT). Significance determined by Tukey’s pairwise comparison.

To verify this interaction is antibody-mediated and not due to a global deficiency in phagocytosis, we also performed a generalized phagocytic uptake assay, wherein BMDM of various genetic backgrounds were cocultured with killed K12 *Escherichia coli* labeled with fluorescein. By pairwise comparison we show that there is no significant difference in phagocytic uptake between DOCF1 mice or CCxCF1 models (**Figure 7D-E**). Representative confocal images of CC024CF1 BMDM uptake of *E. coli* are provided in **Supplement 21**.

We next performed qRT-PCR on a selection of Chr10 locus genes (HACE1, PREP, PRDM1, SNX3), a selection of genes chosen based on differential expression in scRNA-seq data (CTSS, CTSL), and canonical Mφ markers (F4/80, ARG1, NOS2, CCL8). Despite differences observed in ADCP, no significant difference was observed by CCxCF1 model or by response in any of these transcripts (**Supplement 22**). While our genetic analysis did not implicate the F_c_R gene loci (on Chr1 and 3) as genetic drivers, we did find differences in F_c_R mRNA expression in our scRNA-seq data. Interestingly, F_c_R expression is seen primarily in cells capable of ADCP, with almost no detection in NK cells despite their capacity for F_c_R-mediated ADCC (**Supplement 23**). Among phagocytic cells, F_c_gR1 (CD64), F_c_gR2b (CD32b), F_c_gR3 (CD16a) and F_c_gR4 (CD16-2) are differentially expressed between Mφ of different genetic backgrounds, however there is no clear pattern associated with response rate (**Supplement 24A**). Pro-ADCP receptor CD64 is expressed significantly higher in the CR CC031CF1 model than either the low-TDT CC024CF1 model or high-TDT CC035CF1 model (Log_2_FC = 0.28 and 0.27, respectively). Inhibitory receptor CD32b is conversely expressed more highly in the low than high-TDT model (Log_2_FC = 0.19) but not in the CR. Pro-ADCP receptor CD16 is expressed at higher levels in the low versus high-TDT model (Log_2_FC = 0.17), but not in the CR. Lastly, pro-ADCP receptor CD16-2 is expressed at lower levels in the low-than high-TDT model (Log_2_FC = 0.18).

To further investigate whether F_c_R family dysregulation is specific to conditions of the TME, we evaluated expression on *ex vivo*-differentiated BMDM derived from DOCF1 mice and clonal CCxCF1 models by FC (**Supplement 24B**, gating strategy in **Supplement 25**). Similar to the *in vivo* scRNA-seq results, F_c_R expression levels vary but do not correlate to response rate. BMDM from low-TDT DOCF1 mice and CR mice show significantly different expression of CD64, with BMDM from low-TDT tumors expressing ∼1.5x MFI as detected by FC. Additionally, BMDM from low-TDT DOCF1 mice express ∼2x the inhibitory receptor CD32b, and ∼1.5x higher CD16 and CD16-2 expression than all other conditions. No significant differences are seen between BMDM from these clonal models *ex vivo*, suggesting that any F_c_R expression differences between these strains’ Mφ in the TME may be due to signaling that is absent *in vitro*.

## 4 Discussion

Using DOCF1 mice, we find that genetic background predicates response to targeted mAb therapy, and genetic linkage analysis identified a locus on Chr10 (∼40.6-46.6 Mbp) as a candidate driver. PWK genetics at this locus correlate with a positive response to therapy, and conversely A/J correlates to a negative response. We recapitulated this phenotype using three CCxCF1 models and conclude that the Chr10 locus represents a promising lead for investigating regulators of therapeutic anti-tumor antibody response. Of importance, this locus is conserved on human Chr6. *Ex vivo* studies revealed differential capacity for ADCP, but not general phagocytosis, by genotype, suggesting this locus regulates macrophage function with therapeutic anti-tumor antibody.

There is a growing body of evidence that outbred mouse populations may serve as better representative research than traditional clonal models [18], but it is important to note that not all phenotypes can be evaluated in this fashion. A study in 2018 evaluated 107 articles, as well as a JAX DO mouse trait dataset and found that coefficients of variation between inbred and outbred mice only varied significantly in 6/26 metrics considered. This implies that many phenotypes do not vary with genetic background [38]. However, BALB/c mice reliably accept TUBO tumor and consistently behave as partial responders after treatment with mAb clone 7.16.4 immunotherapy, with evidence of CD8^+^ T_c_ cell infiltration with high levels of PD-1 induction [39]. Our study shows that DOCF1 mice widely vary in their response to treatment, supporting the exploration of genetic regulators of this phenotype.

Evaluation of the TME of DOCF1 mice reveals significant elevation of Arg1 and CD8^+^ T_c_ staining in slower-growing tumors. As CD8^+^ T_c_ are not directly engaged by anti-tumor antibody therapy, their increase may imply that antigen presentation is also enhanced in responder mice. Higher Arg1 expression may be surprising given these tumors are more responsive to treatment, but immunosuppressive M2 Mφ and/or MDSC may be induced as a result of smoldering anti-tumor immunity. It is worth remembering that these tumors have radically different harvest dates, so important temporal information regarding the microenvironment may be absent.

Canonical mechanisms of antibody-dependent elimination of tumor tissue include ADCC and ADCP, which are carried out by two major innate immune effector cell populations (NK cells and Mφ respectively). Unsurprisingly, we find that both NK cell and Mφ depletion are detrimental to survivorship after targeted mAb therapy. Bulk RNA seq revealed a core transcriptional program that is deficient in mice who do not respond to therapy. Genes differentially regulated between these two groups implicate ADCP as a core process impacted by the genetics driving this phenotype. While bulk sequencing is informative, the sequencing of select low/high-TDT tumors *post hoc* complicates matters. First, no tumors could be collected from true CR mice; second, due to differences in harvest dates our results are subject to temporal flux in the immune response.

To obtain a clearer picture of the TME during the therapeutic inflection point, prior to CR mice eliminating tumor, scRNA-seq was performed in our CCxCF1 models. The immune fractions of all strains are dominated by various subsets of Mφ and T cells, with additional smaller populations of monocytes, NK cells, neutrophils, cDC, pDC, and B cells. Though not explorde further here, we found that neutrophils are more abundant in low-TDT tumors as compared to responder strains. It has been shown that neutrophils contribute to tumor progression through multiple mechanisms, including suppression of the adaptive immune response by expression of arginase (ARG1), reactive oxygen species (ROS), and nitric oxide (NO) in the TME [40]. This is further evidenced by the predictive value of neutrophil to lymphocyte ratios in the context of neoadjuvant HER2-targeted therapies trastuzumab and pertuzumab [41]. Other processes that implicate neutrophils as negative regulators of the immunotherapy response include the promotion of genetic instability, tumor cell proliferation, angiogenesis, and metastasis (reviewed in [42]). It should be noted that polymorphonuclear myeloid-derived suppressor cells (PMN-MDSC) share developmental origins, markers, and many functional features with neutrophils. Although these cells have been identified by SingleR as neutrophil, the possibility remains that these may be better categorized as MDSC and further investigation is warranted.

In addition to monocytes, DC, and Mφ subset overrepresentation in high-TDT/CR tumors, we found NK cells are also overrepresented in high-TDT/CR models. NK cells are known to contribute to anti-tumor immunity, but in many human cancers NK cells are only present in low numbers [12]. B cells were also more abundant in high-TDT/CR strains, and although they are the least populous subset of the immune infiltrate, B cells have previously been reported to impact response to targeted immunotherapies. Generating host-derived HER2-specific antibody is beneficial to patients, both for reducing relapse among patients that have undergone HER2^+^ tumor resection [43] and increasing survival among patients treated with trastuzumab and chemotherapeutics [44].

Mφ have long been implicated as mediators of tumor cell killing during mAb therapy, and it has also been shown that phagocytosis is induced during trastuzumab treatment in breast cancer [45], as well as other cancer models [46]. Mφ in the TME may conversely promote tumor growth through various mechanisms, including the secretion of anti-inflammatory cytokines (TGFβ, IL-10), as well as checkpoint molecule PD-L1, and Arg1 which actively inhibits T-cell activation [47]. Subsetting the Mφ population in our scRNA-seq data reveals 4 subpopulations that distinguish themselves from the core transcriptional Mφ program. We see two key populations overrepresented in high-TDT/CR tumor infiltrate. The first overrepresented transcriptional state expresses genes associated with proinflammatory processes and neutrophil recruitment. This is largely reflected in the targeted immunotherapy literature, as inflammatory Mφ in the M1 phenotype have been associated with increased survival [48]. Our results differ, however, as we see increased levels of neutrophils present in our low-TDT model. The second Mφ subset found overrepresented in high-TDT/CR tumors expresses genes known to regulate endocytosis and monocyte recruitment. Given the critical role ADCP is playing in the targeted immunotherapy response, it is unsurprising that genes involved in endocytosis are upregulated in models that respond well to therapy.

We show that BMDM generated from DOCF1 mice that eliminate tumor more efficiently phagocytose tumor in an Ab-dependent manner *ex vivo*. Importantly, we show that this observation holds true in alternative models of targeted mAb therapy (e.g. B16F0 tumor with anti-TYRP1 antibody), indicating that this observation may be more broadly applicable across multiple tumor types. No significant differences were detected in Ab-independent phagocytosis between groups, further suggesting that genetics are regulating an Ab-dependent process.

While the F_c_R loci were not implicated in our genetic scan, F_c_R dependence is key to the impact of targeted immunotherapy. In our models, F_c_R mediated killing seems to be predominately the domain of intratumoral Mφ. We were surprised to detect virtually no F_c_R mRNA expression in the NK cell population, although it has been shown previously that F_c_R signaling is dispensable for NK cells in the response to targeted immunotherapy [17]. This finding in our dataset might suggest that phagocytic cells are playing a bigger role than NK cell in F_c_R dependent killing at this timepoint in these models.

Some differences were observed in F_c_R expression in our scRNA-seq data. Pro-ADCP receptors CD64 and CD16-2 are expressed at a slightly higher level in high-TDT/CR strains, although the expression patterns are not consistent between responder strains. Further, when unstimulated BMDM from both the diverse and clonal models were tested for F_c_R expression at the protein level *ex vivo*, we see that the TME may be necessary for this phenotype, with the exception of low-TDT DOCF1 mice. This differential regulation may be due to factors within the TME, as many signaling molecules are known modulate F_c_R expression [49,50].

While the processes leveraged to induce immune-mediated killing have been well-described, the complexity of the immune system has made it difficult to tease apart the mechanisms by which these therapies function or fail *in vivo*. Diverse genetic models represent an important tool in the repertoire of scientists seeking to find loci driving complex phenotypes. Here we identify a locus on the murine Chr10 that associates with targeted immunotherapy response, and more specifically with the capacity for Mφ to perform ADCP. This function is core to the cellular identity of tumor infiltrating Mφ and represents a key process that should be seriously considered as a target for modulation in the context of targeted immunotherapeutics. Given the phagocytic capacity of Mφ, their abundance in the TME, and the plasticity of the cell type, the possibility for these cells to be manipulated to the benefit of the patient remains a promising area of research.

## Acknowledgements

We thank Professor Wei-Zen Wei for her contribution of the TUBO cell line, the DLAR at WSU for their care of research subjects involved, the WSU GSC for data processing, and the MICR Core at WSU for their guidance in microscopy, imaging, and flow cytometry.

