## Supplemental Figures for "Host genetic background regulates the capacity for anti-tumor antibody-dependent phagocytosis"

| Primer | Species |
| --- | --- |
| Gapdh (Mm99999915_g1) | mouse |
| B2m (Mm00437762_m1) | mouse |
| Ccl8 (Mm01297183_m1) | mouse |
| Ctsl (Mm00515597_m1) | mouse |
| Ctss (Mm01255859_m1) | mouse |
| Prdm1 (Mm00476128_m1) | mouse |
| Snx3 (Mm01181342_m1) | mouse |
| Hace1 (Mm00552702_m1) | mouse |
| Prep (Mm00448377_m1) | mouse |
| Adgre1 (F4/80) (Mm00802529_m1) | mouse |
| Nos2 (Mm00440502_m1) | mouse |
| Arg1 (Mm00475988_m1) | mouse |
| Neu (Rn00566561_m1) | rat |

***Supplemental Table 1*****. qRT-PCR primers.**

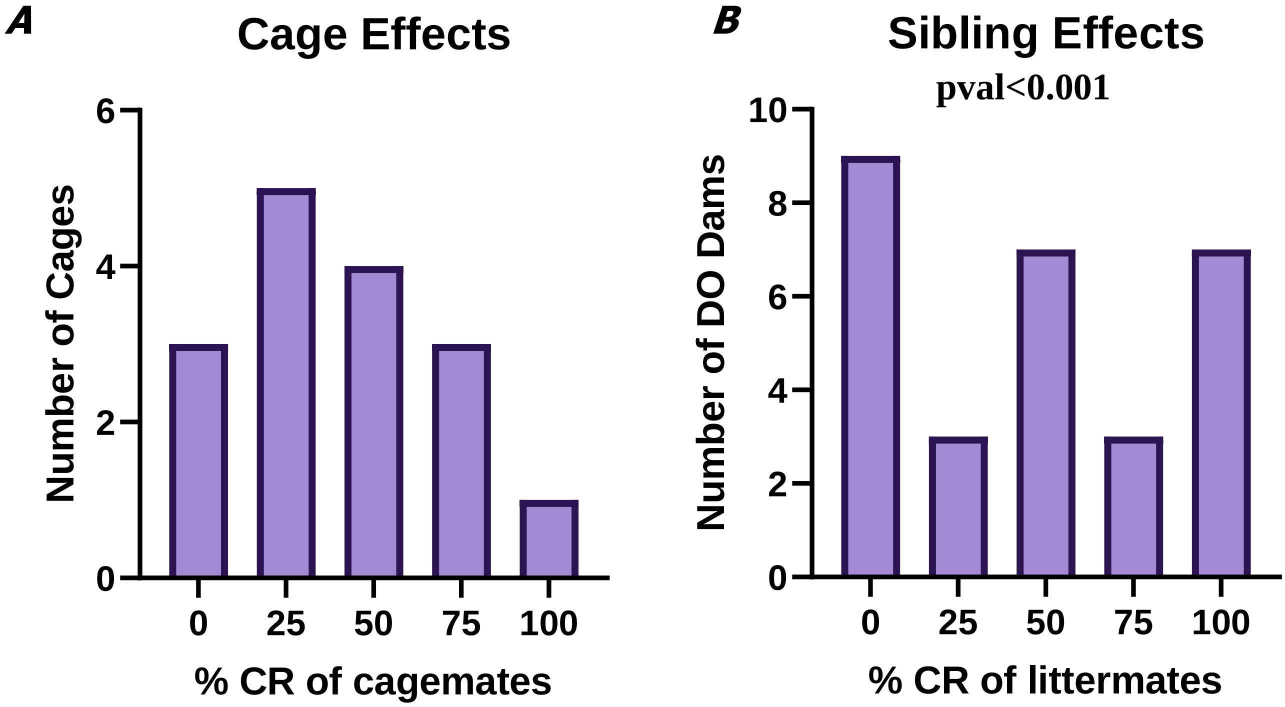


***Supplemental Figure 1.*** **Demonstration of phenotypic heritability of response to targeted immunotherapy in DOCF1 mice.** *(A)* Gaussian distribution of the impact of cohabitation on response to mAb treatment. ***(B)*** Skewed distribution of kinship as a predictor of robust response to mAb treatment.

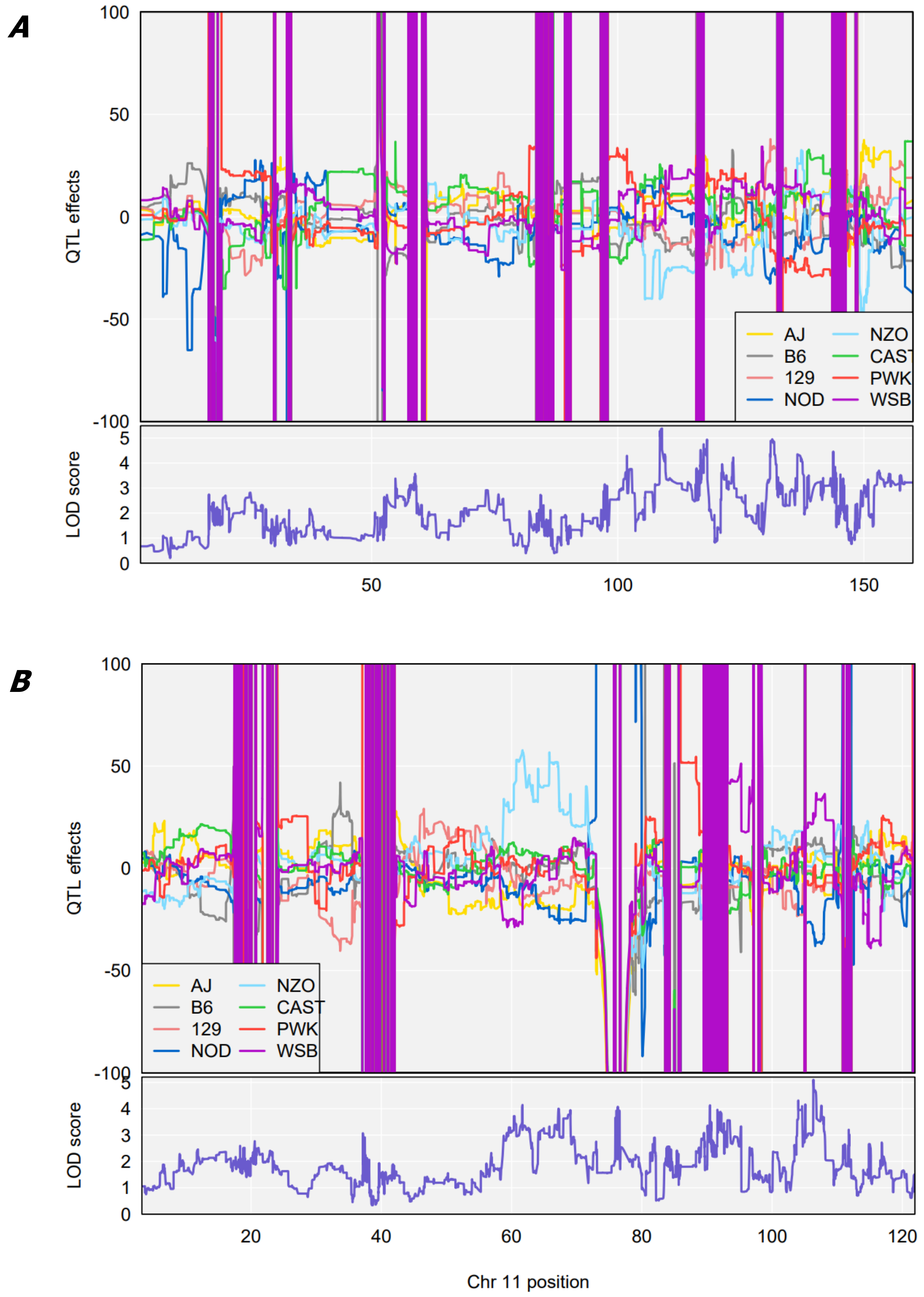


***Supplemental Figure 2.*** **Genetic Linkage Analysis of alternative loci.** ***(A)*** Position of interest on Chr 3, Above: Quantitative Trait Loci (QTL) effect plot at peak. Below: Corresponding LOD scores. ***(B)*** Position of interest on Chr 11, Above: Quantitative Trait Loci (QTL) effect plot at peak. Below: Corresponding LOD scores.

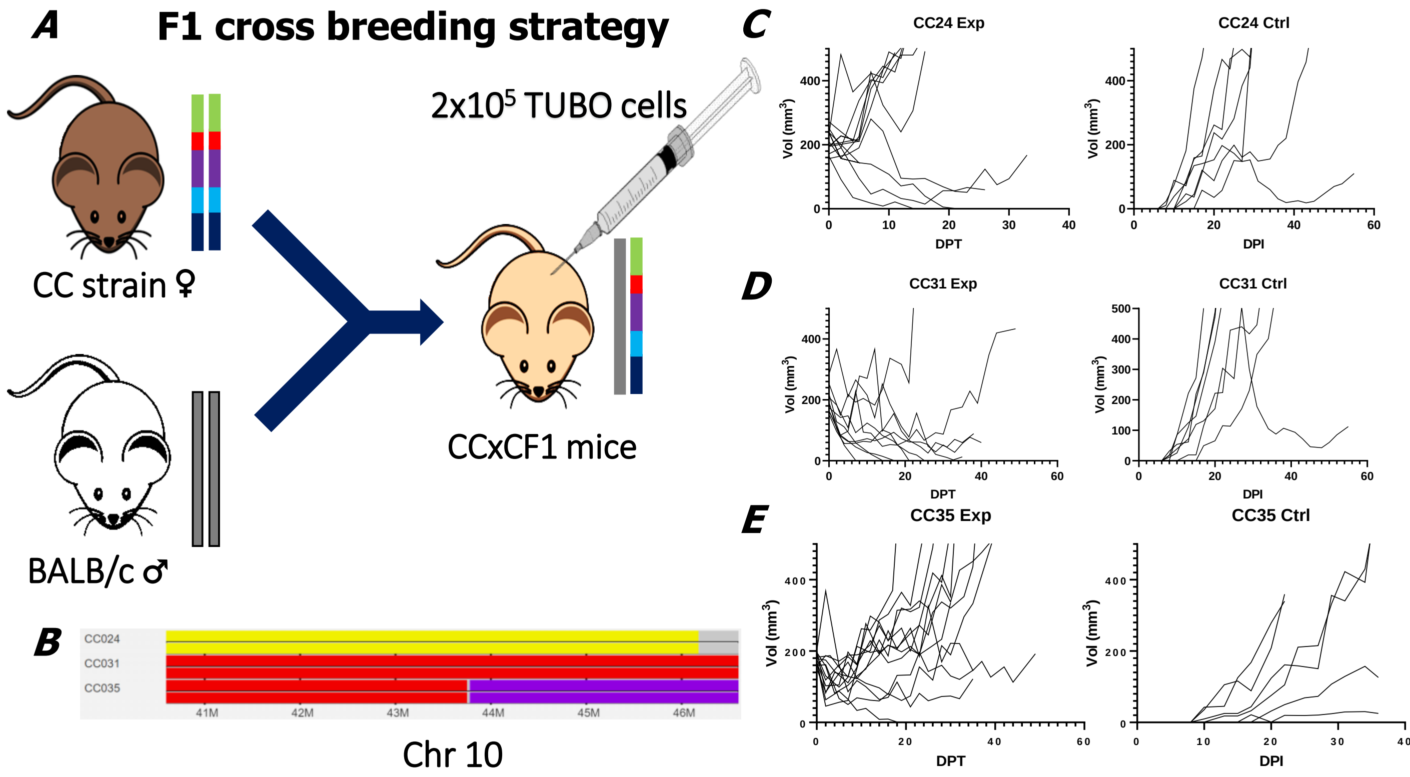


***Supplemental Figure 3.*** **Collaborative Cross F1 cross experimental design and tumor growth curves.** ***(A)*** Breeding scheme for development of CCxCF1 mice, capable of readily accepting orthotopic implantation of TUBO. ***(B)*** Founder strain contribution at locus of interest. ***(C)*** CC024CF1 tumor growth curves, treated (Exp) vs untreated (Ctrl) ***(D)*** CC031CF1 tumor growth. ***(E)*** CC035CF1 tumor growth.

**A**

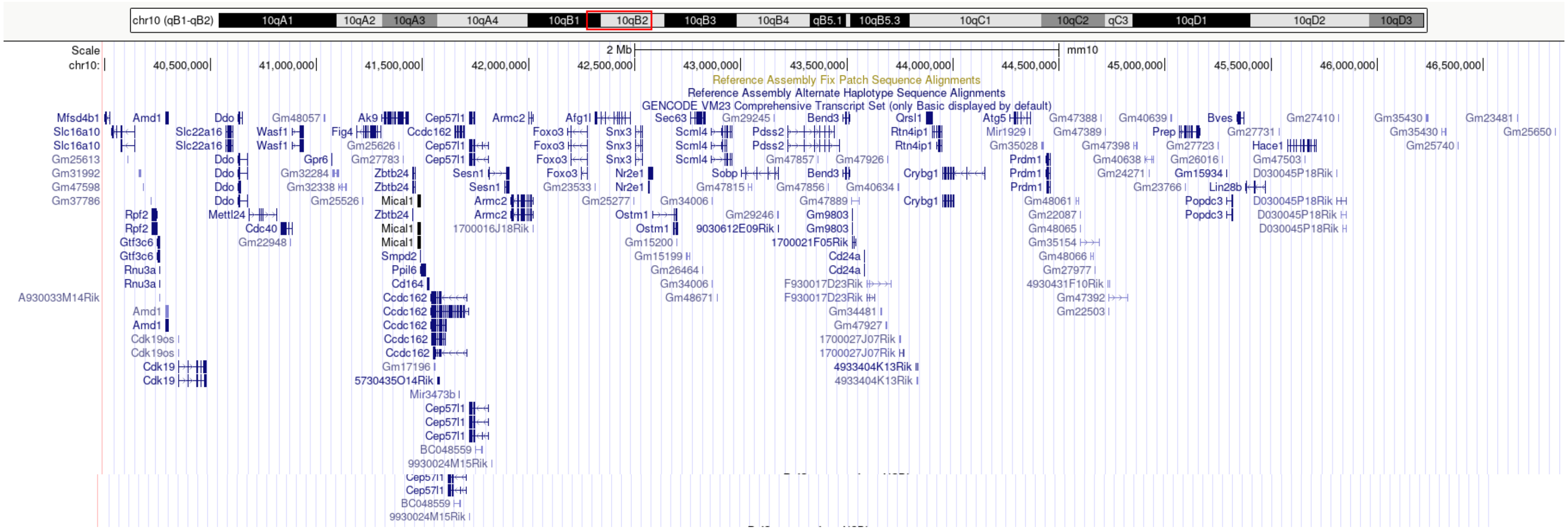

**B**

| OFFICIAL_GENE_SYMBOL | GENE NAME |
|---|---|
| Bend3 | BEN domain containing 3(Bend3) |
| Hace1 | HECT domain and ankyrin repeat containing, E3 ubiquitin protein ligase 1(Hace1) |
| Prdm1 | PR domain containing 1, with ZNF domain(Prdm1) |
| Scml4 | Scm polycomb group protein like 4(Scml4) |
| Foxo3 | forkhead box O3(Foxo3) |
| Mtres1 | mitochondrial transcription rescue factor 1(Mtres1) |
| Nr2e1 | nuclear receptor subfamily 2, group E, member 1(Nr2e1) |

***Supplemental Figure 4.*** **Genetic map of locus of interest.** ***(A)*** Map of Chr 10 40-47Mbp. ***(B)*** Transcription Factors present at this locus.

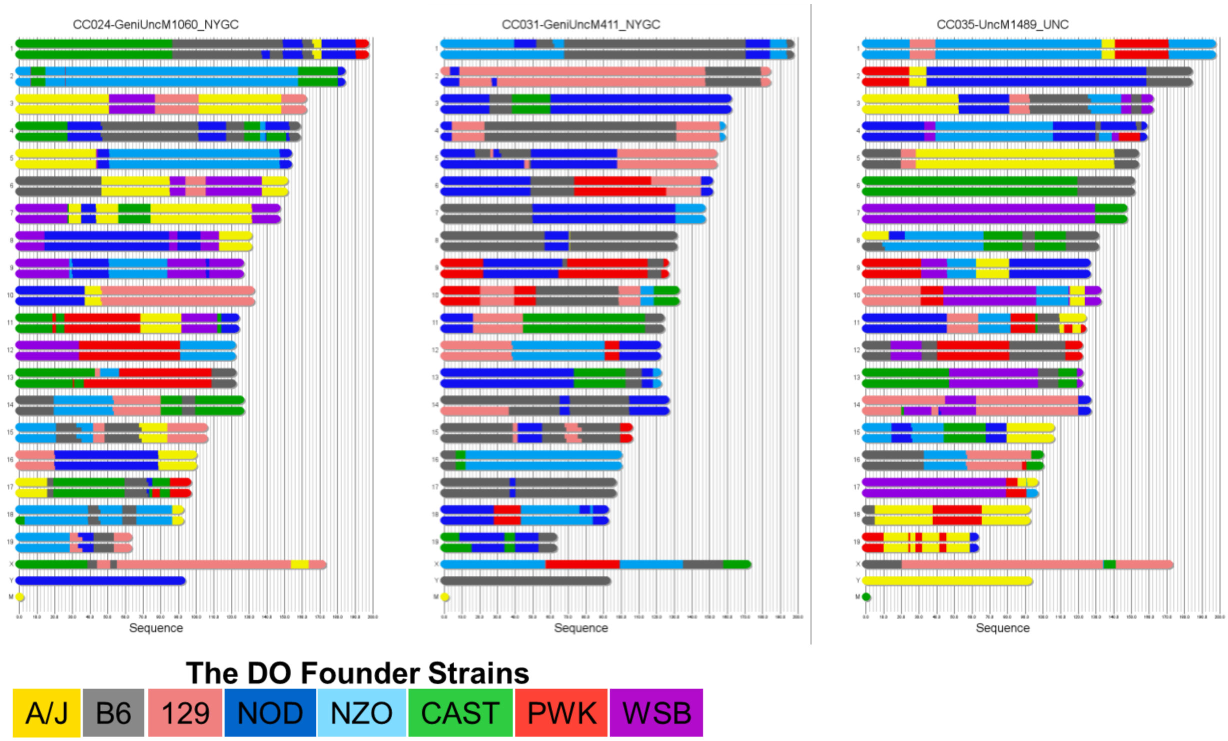


***Supplemental Figure 5.*** **CC founder genomes.** Founder contributions across the relevant CC genomes (CC024, CC031, CC035) and color scheme representing the 8 founder genomes comprising the Diversity Outbred mouse model.

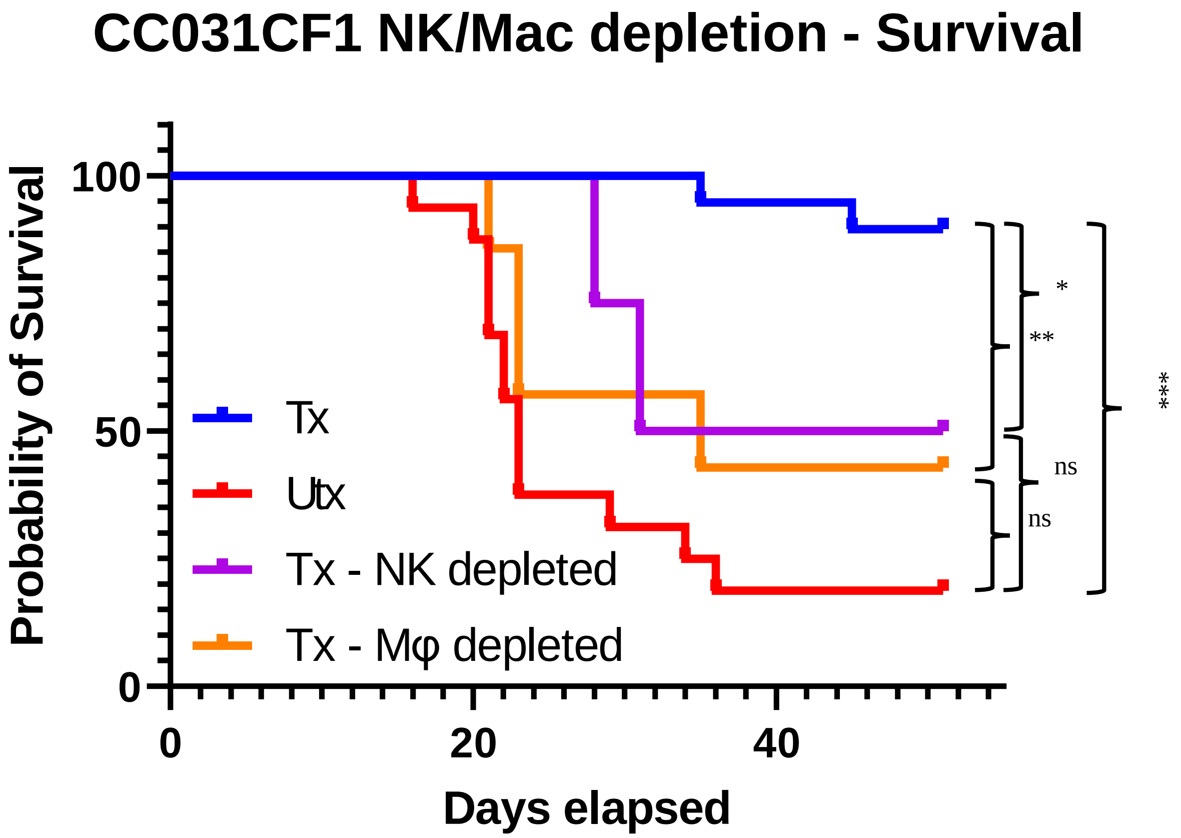


***Supplemental Figure 6.* Depletion of key innate immune cell populations negatively impacts survival of CR model CC031CF1 during targeted immunotherapy treatment.** Kaplan–Meier curve depicting survivorship of 7.16.4-treated TUBO-bearing CC031CF1 mice with and without NK cell (anti-AsGM1) or Mφ (anti-Csf1r) depletion. Historical Tx and Utx mice included.

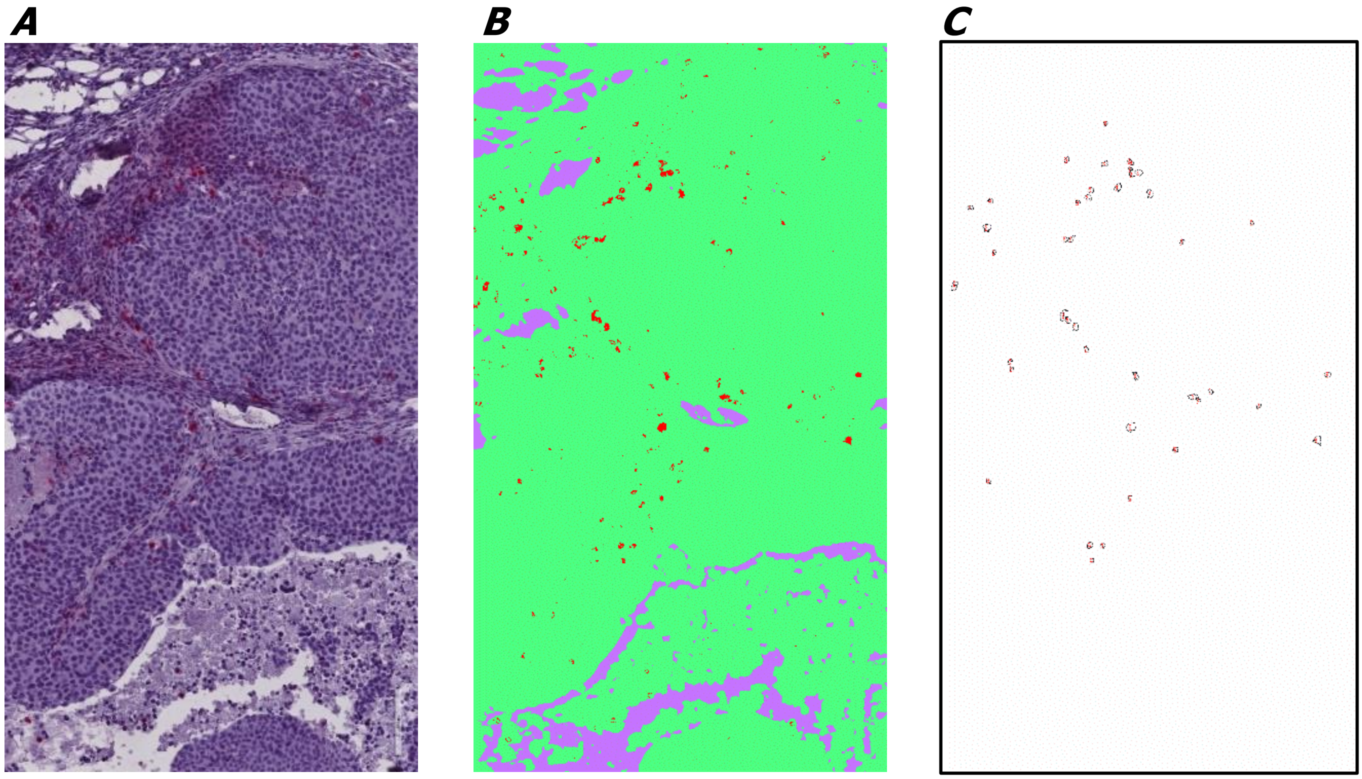


***Supplemental Figure 7.*** **Strategy for quantification of immune infiltrate in IHC stained tumor tissue.** ***(A)*** Original image. ***(B)*** Post-Trainable Weka segmentation. ***(C)*** Quantification of positive cells.

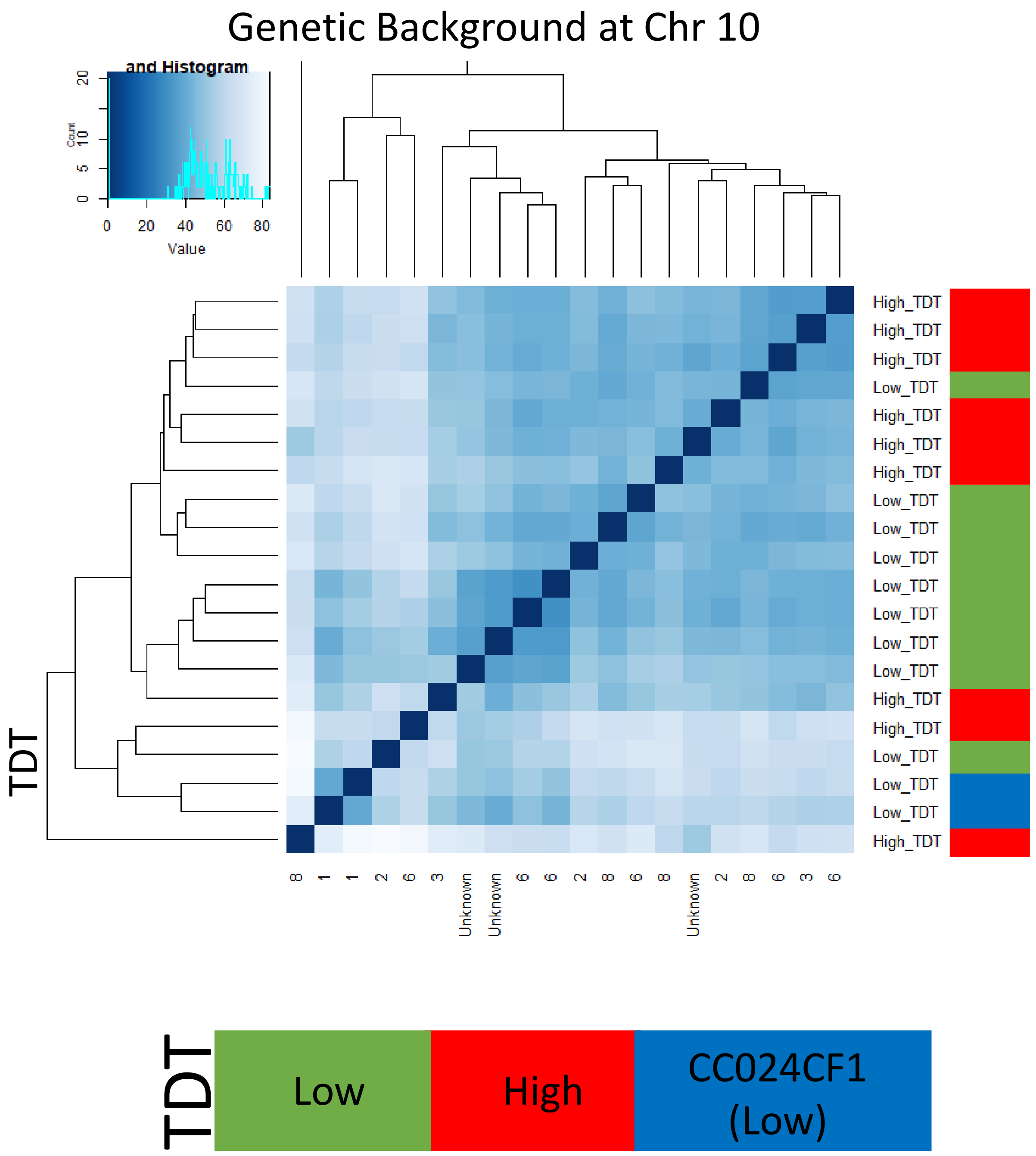


***Supplemental Figure 8.*** **QuantSeq transcriptomic analysis of flash frozen tumor**. 8 High DOCF1 TDT samples, 8 Low TDT DOCF1 samples, and two CC024C. Distance matrix on based on the relatedness of each sample's transcriptome.

| Annotation Cluster 1 | Enrichment Score: 6.61 | G | | Count | P_Value | Benjamini |
|---|---|---|---|---|---|---|
| GOTERM_BP_DIRECT | immune system process | RT | | 63 | 3.4E-13 | 1.4E-9 |
| GOTERM_BP_DIRECT | defense response to virus | RT | | 32 | 2.7E-8 | 3.7E-5 |
| UP_KW_BIOLOGICAL_PROCESS | Innate immunity | RT | | 41 | 2.4E-7 | 1.4E-5 |
| UP_KW_BIOLOGICAL_PROCESS | Immunity | RT | | 63 | 4.2E-7 | 1.7E-5 |
| UP_KW_BIOLOGICAL_PROCESS | Antiviral defense | RT | | 18 | 2.9E-6 | 8.8E-5 |
| GOTERM_BP_DIRECT | response to virus | RT | | 16 | 1.8E-5 | 9.3E-3 |
| GOTERM_BP_DIRECT | innate immune response | RT | | 49 | 1.1E-3 | 1.7E-1 |
| **Annotation Cluster 2** | **Enrichment Score: 3.08** | **G** | | **Count** | **P_Value** | **Benjamini** |
| KEGG_PATHWAY | Epstein-Barr virus infection | RT | | 27 | 5.8E-5 | 3.5E-3 |
| KEGG_PATHWAY | Cellular senescence | RT | | 23 | 8.9E-5 | 3.5E-3 |
| KEGG_PATHWAY | Human immunodeficiency virus 1 infection | RT | | 24 | 1.5E-3 | 2.3E-2 |
| KEGG_PATHWAY | Human cytomegalovirus infection | RT | | 25 | 1.6E-3 | 2.3E-2 |
| KEGG_PATHWAY | Kaposi sarcoma-associated herpesvirus infection | RT | | 22 | 3.1E-3 | 3.4E-2 |
| KEGG_PATHWAY | Viral carcinogenesis | RT | | 21 | 8.4E-3 | 7.8E-2 |
| **Annotation Cluster 3** | **Enrichment Score: 2.69** | **G** | | **Count** | **P_Value** | **Benjamini** |
| GOTERM_CC_DIRECT | late endosome membrane | RT | | 17 | 8.9E-5 | 6.0E-3 |
| GOTERM_CC_DIRECT | lysosomal membrane | RT | | 26 | 1.2E-4 | 7.4E-3 |
| GOTERM_CC_DIRECT | lysosome | RT | | 29 | 3.3E-2 | 3.5E-1 |
| UP_KW_CELLULAR_COMPONENT | Lysosome | RT | | 24 | 4.6E-2 | 2.1E-1 |
| **Annotation Cluster 4** | **Enrichment Score: 2.45** | **G** | | **Count** | **P_Value** | **Benjamini** |
| GOTERM_BP_DIRECT | proteasome-mediated ubiquitin-dependent protein catabolic process | RT | | 22 | 4.0E-4 | 9.0E-2 |
| GOTERM_CC_DIRECT | proteasome core complex, beta-subunit complex | RT | | 5 | 7.9E-4 | 2.3E-2 |
| GOTERM_CC_DIRECT | proteasome core complex | RT | | 6 | 9.0E-4 | 2.5E-2 |
| GOTERM_BP_DIRECT | proteasomal ubiquitin-independent protein catabolic process | RT | | 6 | 2.0E-3 | 2.4E-1 |
| GOTERM_BP_DIRECT | proteolysis involved in cellular protein catabolic process | RT | | 10 | 2.0E-3 | 2.4E-1 |

***Supplemental Figure 9.*** **Gene Ontology functional annotation clustering.** Functional enrichment of differentially expressed genes in High vs Low TDT via Quantseq transcriptome analysis. Overrepresented GO terms: Biological process, Pathway, Cellular Component ontologies. Analysis by https://david.ncifcrf.gov/.

| ID | GENE NAME |
|---|---|
| ATPase, class V, type 10B(Atp10b) | ATPase, class V, type 10B(Atp10b) |
| CD74 antigen (invariant polypeptide of major histocompatibility complex, class II antigen-associated)(Cd74) | CD74 antigen (invariant polypeptide of major histocompatibility complex, class II antigen-associated)(Cd74) |
| SID1 transmembrane family, member 2(Sidt2) | SID1 transmembrane family, member 2(Sidt2) |
| WD repeat domain 81(Wdr81) | WD repeat domain 81(Wdr81) |
| aldo-keto reductase family 1, member B10 (aldose reductase)(Akr1b10) | aldo-keto reductase family 1, member B10 (aldose reductase)(Akr1b10) |
| ankyrin 3, epithelial(Ank3) | ankyrin 3, epithelial(Ank3) |
| cathepsin A(Ctsa) | cathepsin A(Ctsa) |
| cathepsin H(Ctsh) | cathepsin H(Ctsh) |
| cytotoxic T lymphocyte-associated protein 2 alpha(Ctla2a) | cytotoxic T lymphocyte-associated protein 2 alpha(Ctla2a) |
| deoxyribonuclease II alpha(Dnase2a) | deoxyribonuclease II alpha(Dnase2a) |
| dynein cytoplasmic 1 light intermediate chain 1(Dync1li1) | dynein cytoplasmic 1 light intermediate chain 1(Dync1li1) |
| dynein, cytoplasmic 1 light intermediate chain 2(Dync1li2) | dynein, cytoplasmic 1 light intermediate chain 2(Dync1li2) |
| histocompatibility 2, class II, locus Mb1(H2-DMb1) | histocompatibility 2, class II, locus Mb1(H2-DMb1) |
| immunity-related GTPase family M member 1(Irgm1) | immunity-related GTPase family M member 1(Irgm1) |
| interferon gamma inducible protein 30(Ifi30) | interferon gamma inducible protein 30(Ifi30) |
| interferon induced transmembrane protein 3(Ifitm3) | interferon induced transmembrane protein 3(Ifitm3) |
| kinesin family member 5B(Kif5b) | kinesin family member 5B(Kif5b) |
| late endosomal/lysosomal adaptor, MAPK and MTOR activator 4(Lamtor4) | late endosomal/lysosomal adaptor, MAPK and MTOR activator 4(Lamtor4) |
| late endosomal/lysosomal adaptor, MAPK and MTOR activator 5(Lamtor5) | late endosomal/lysosomal adaptor, MAPK and MTOR activator 5(Lamtor5) |
| low density lipoprotein receptor(Ldlr) | low density lipoprotein receptor(Ldlr) |
| metallothionein 1(Mt1) | metallothionein 1(Mt1) |
| ring finger protein 152(Rnf152) | ring finger protein 152(Rnf152) |
| solute carrier family 29 (nucleoside transporters), member 3(Slc29a3) | solute carrier family 29 (nucleoside transporters), member 3(Slc29a3) |
| sorting nexin 14(Snx14) | sorting nexin 14(Snx14) |
| transient receptor potential cation channel, subfamily M, member 2(Trpm2) | transient receptor potential cation channel, subfamily M, member 2(Trpm2) |
| transmembrane protein 165(Tmem165) | transmembrane protein 165(Tmem165) |
| unc-93 homolog B1, TLR signaling regulator(Unc93b1) | unc-93 homolog B1, TLR signaling regulator(Unc93b1) |
| vesicle transport through interaction with t-SNAREs 1B(Vti1b) | vesicle transport through interaction with t-SNAREs 1B(Vti1b) |
| vesicle-associated membrane protein 8(Vamp8) | vesicle-associated membrane protein 8(Vamp8) |

***Supplemental Figure 10.*** **Gene Ontology functional annotation clustering.** DEG via Quantseq transcriptome analysis, GO terms: Lysosomal Genes. Analysis by https://david.ncifcrf.gov/.

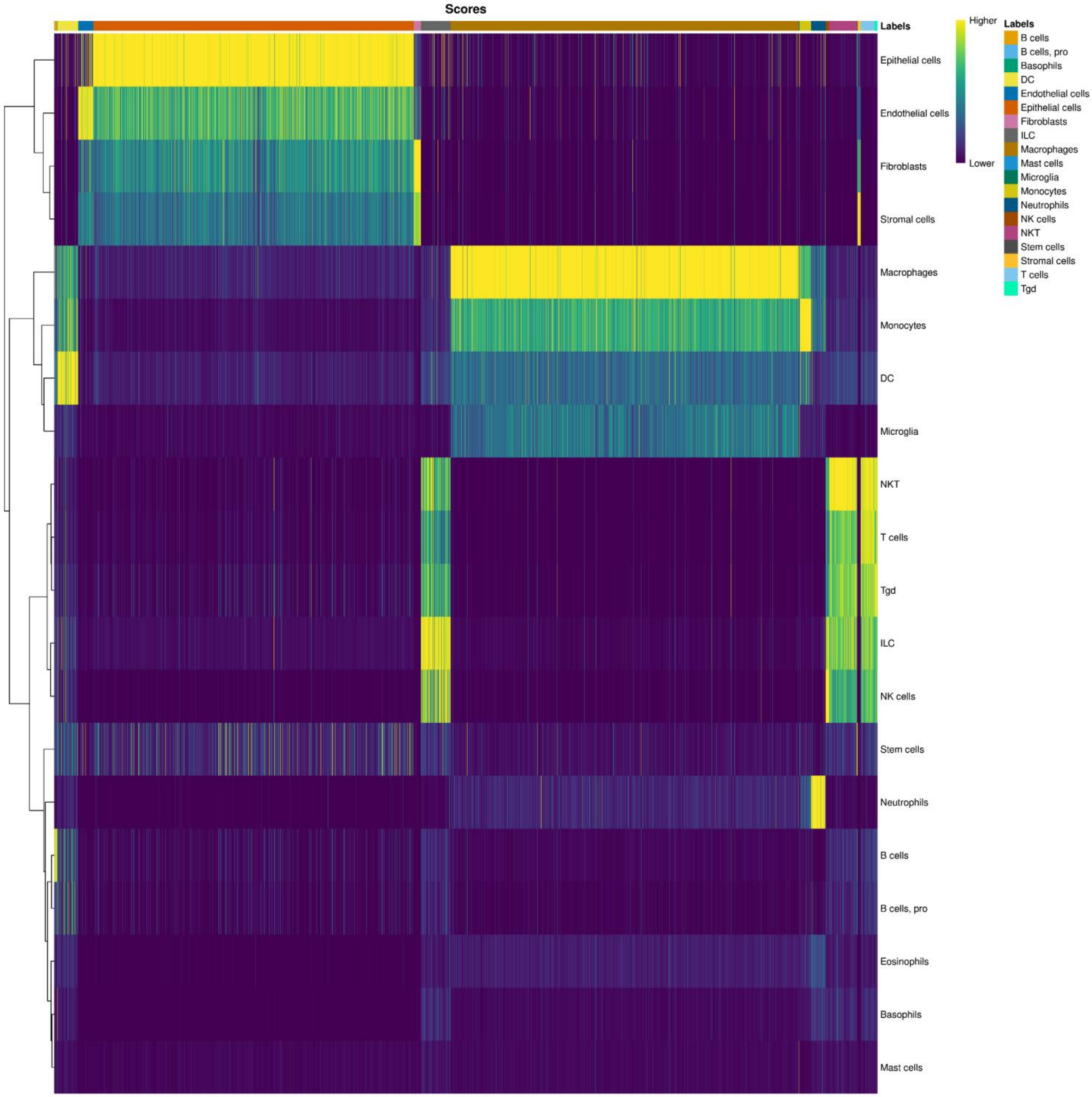


***Supplemental Figure 11.* SingleR identification of single cells.** Individual Spearman Correlation scores assigning the probability that each cell is likely each of the given listed immune cell subsets.

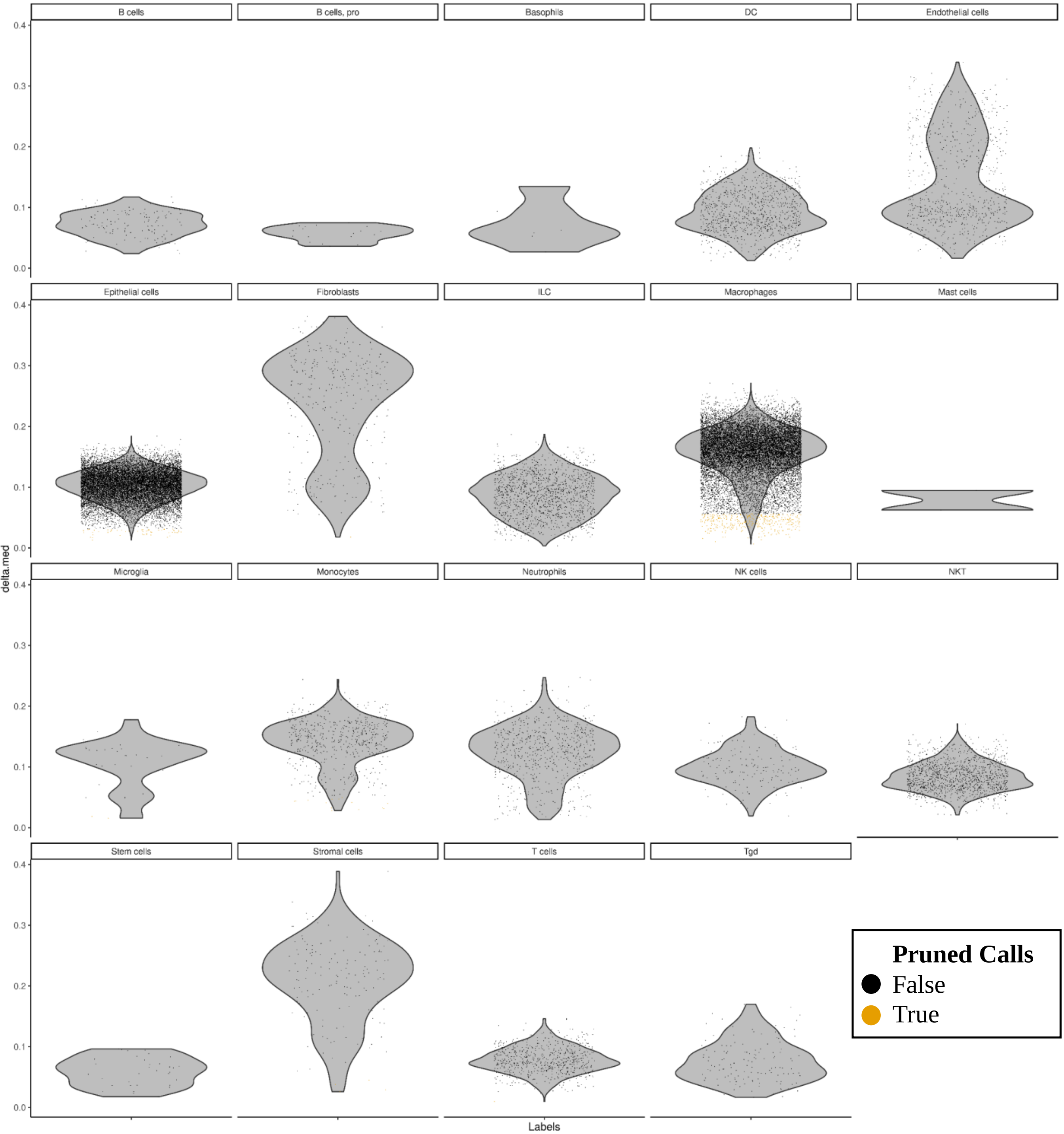


***Supplemental Figure 12.*** **SingleR identification of single cells.** Deviance from each cells assigned ID and the mean spearman correlation score (delta score).

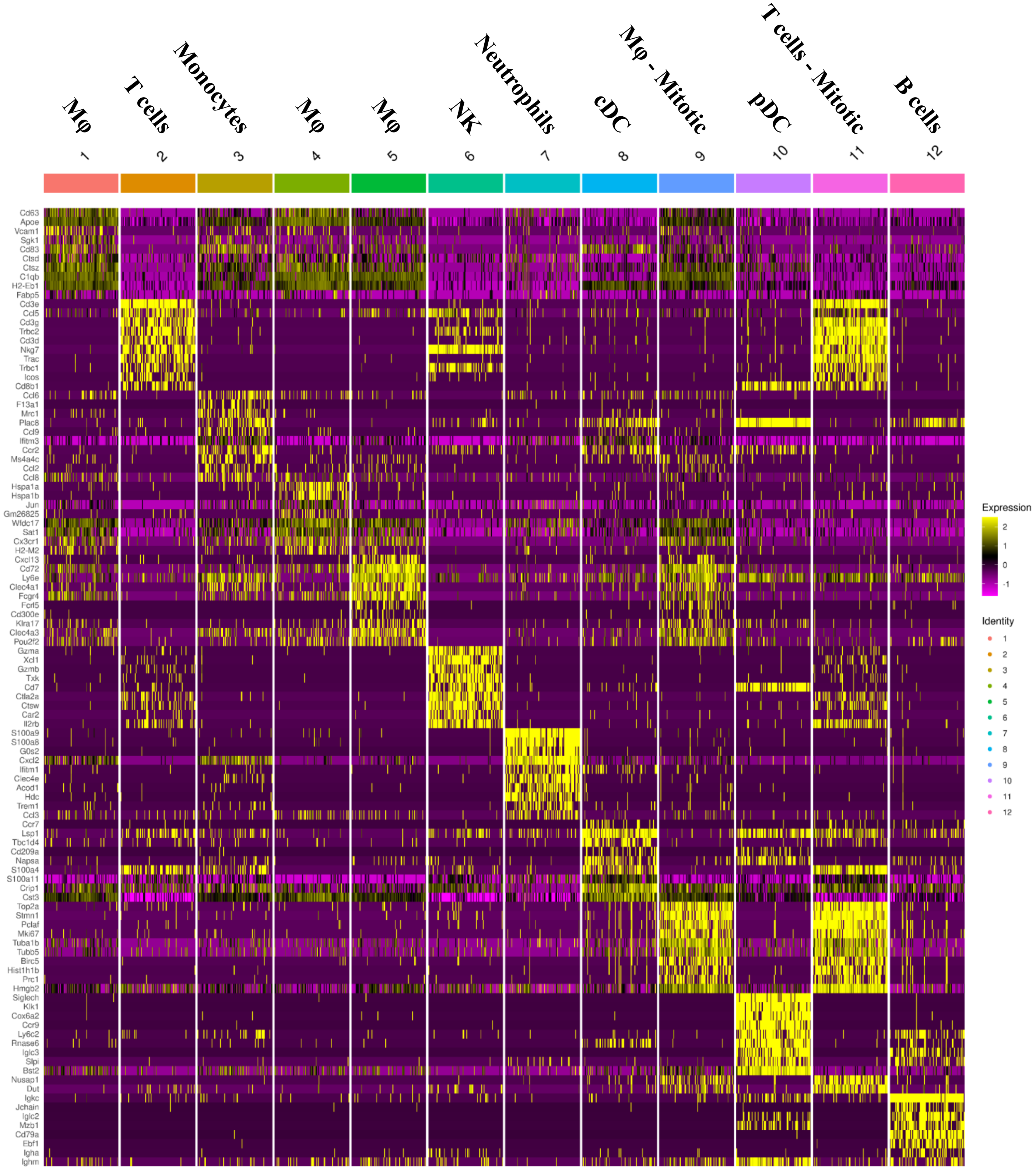


***Supplemental Figure 13.*** **Transcriptional signatures of CD45$^+$ tumor infiltrate.** Top 10 differentially expressed genes across CD45$^+$ clusters.

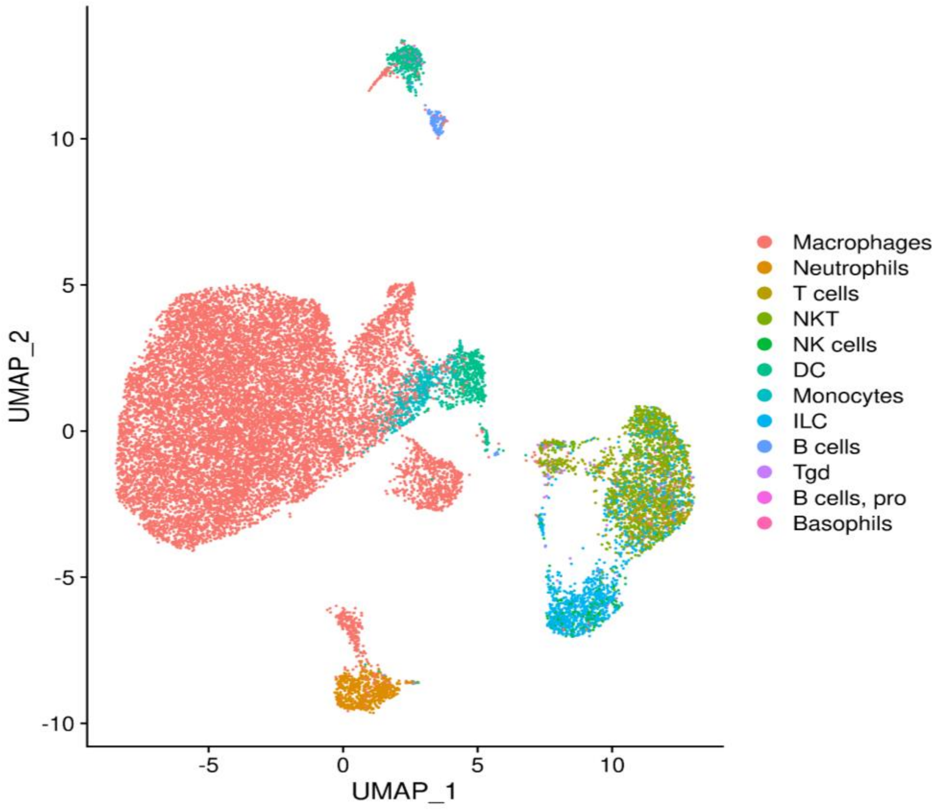


***Supplemental Figure 14.*** **SingleR identification of single cells.** $CD45^+$ fraction.

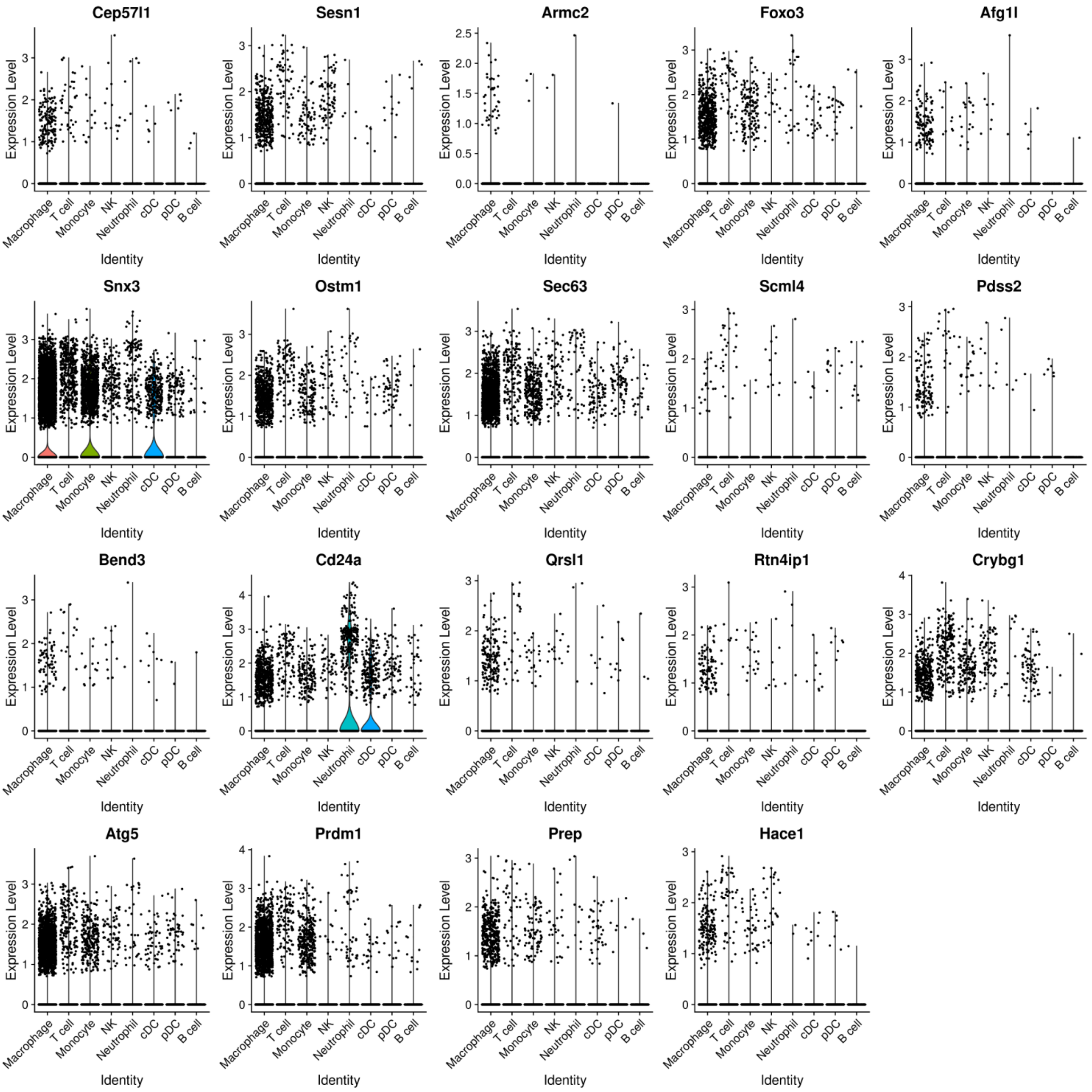


***Supplemental Figure 15.*** **scRNA identifies Chr 10 locus expression.** Transcription of protein-coding genes in the Chr 10 locus of interest, by identified cell types.

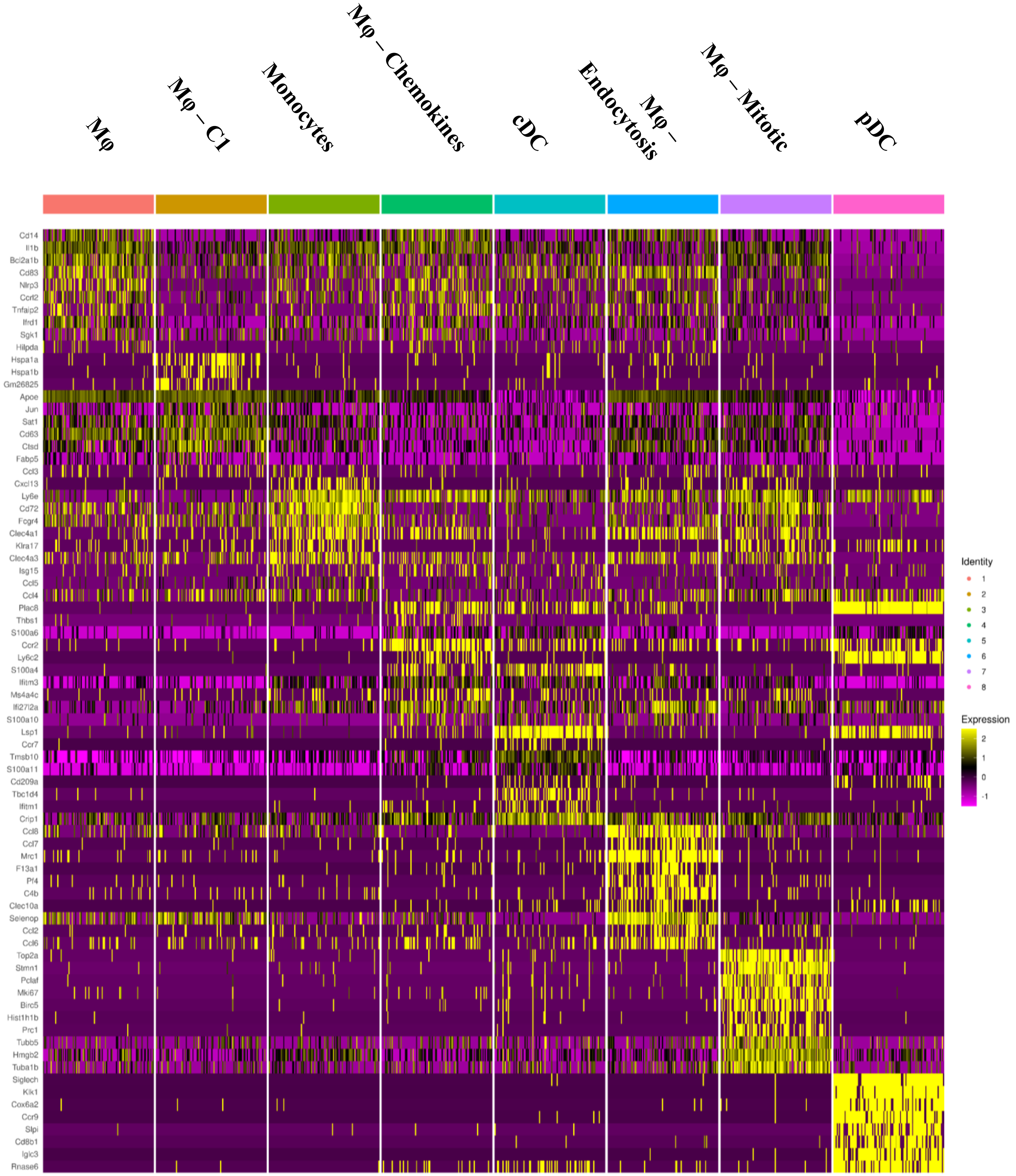


***Supplemental Figure 16.*** **Transcriptional signatures subclustered infiltrating monocytes, macrophage, and DCs.** Top 10 differentially expressed genes across clusters.

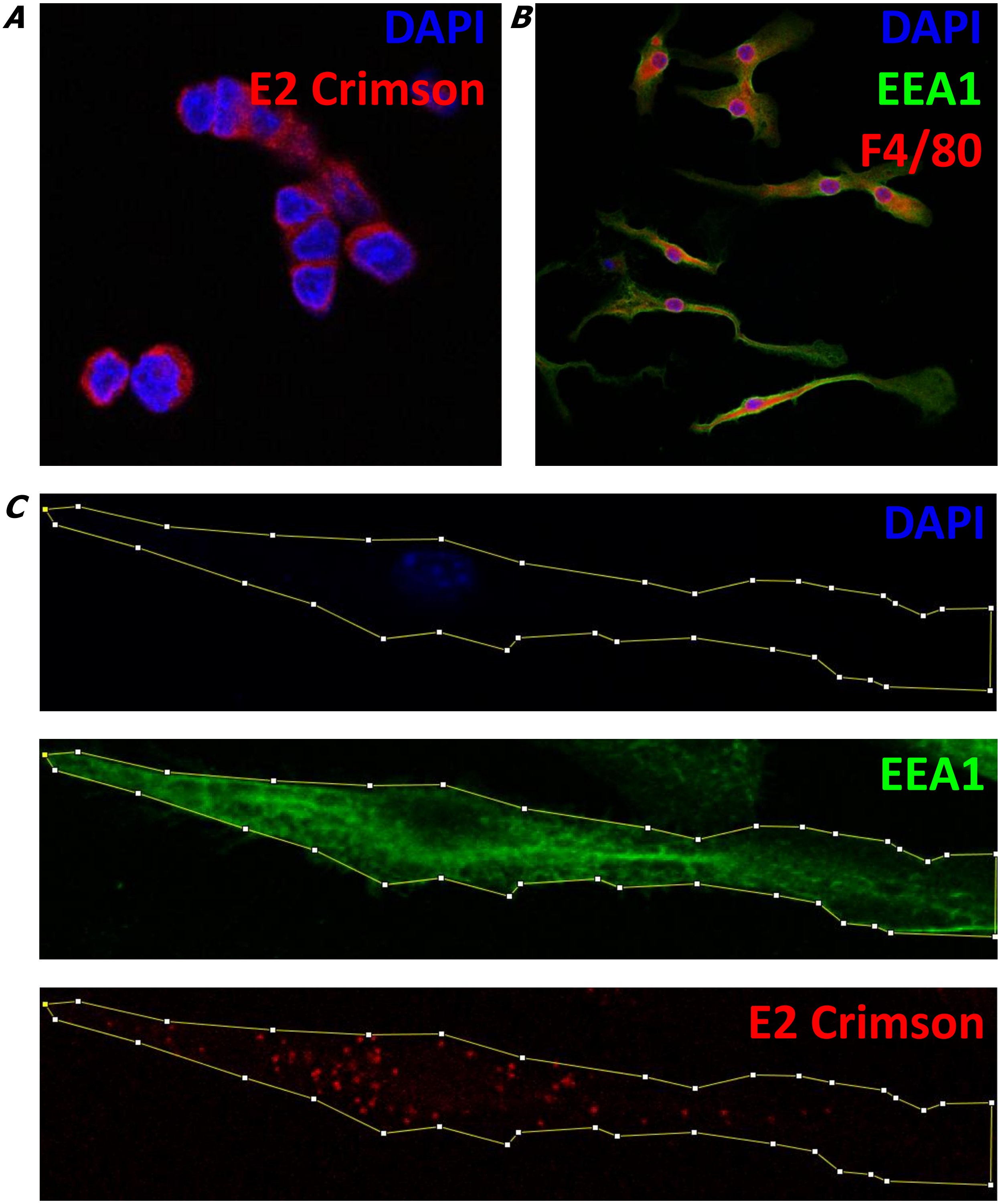


***Supplemental Figure 17.*** **Confirmation of Macrophage differentiation and ADCP quantification.** ***(A)*** TUBO-crimson monoculture, stained for DAPI (blue), and expressing E2-Crimson (red). ***(B)*** BMDM derived in culture, 7 days post seeding in 30% L-929 supplemented R10 media, stained for F4/80 (red), DAPI (blue), and EEA1 (green). ***(C)*** Method used to calculate Mean Fluorescent Intensity (MFI) of phagocytosed E2-Crimson via FIJI (FIJI is just imageJ).

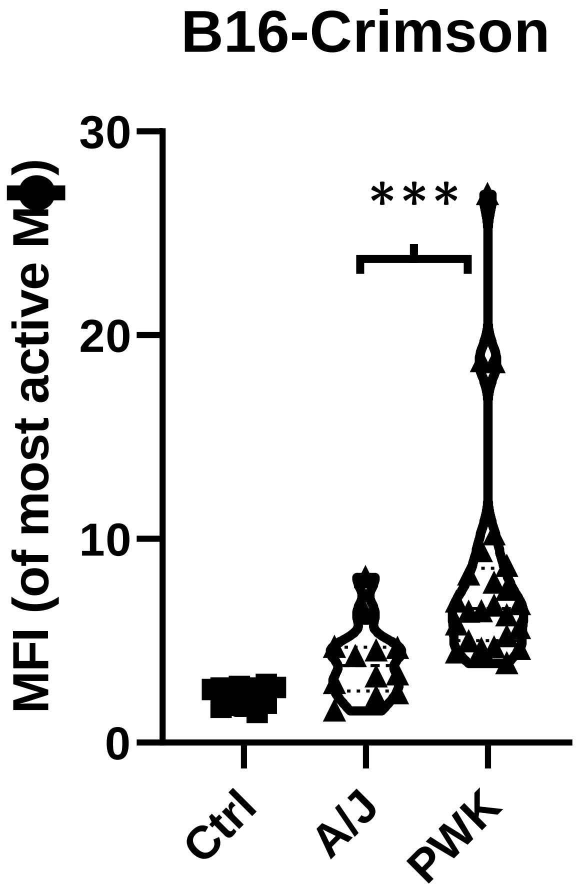


***Supplemental Figure 18.*** **Genetic background in CCxCF1 BMDM predicates higher rates of ADCP ex vivo in alternative model systems.** Bone Marrow Derived Macrophage (BMDM) derived from CC024CF1 (A/J), CC031CF1 (PWK), and CC035CF1 (PWK) in coculture with opsonized B16/Crimson tumor cells (mAb BE0151 – anti-TA99) for 4 hours. MFI of representative BMDM, selected across three fields of view. Control (Ctrl) – Non-opsonized, analysis by Mann Whitney t-test.

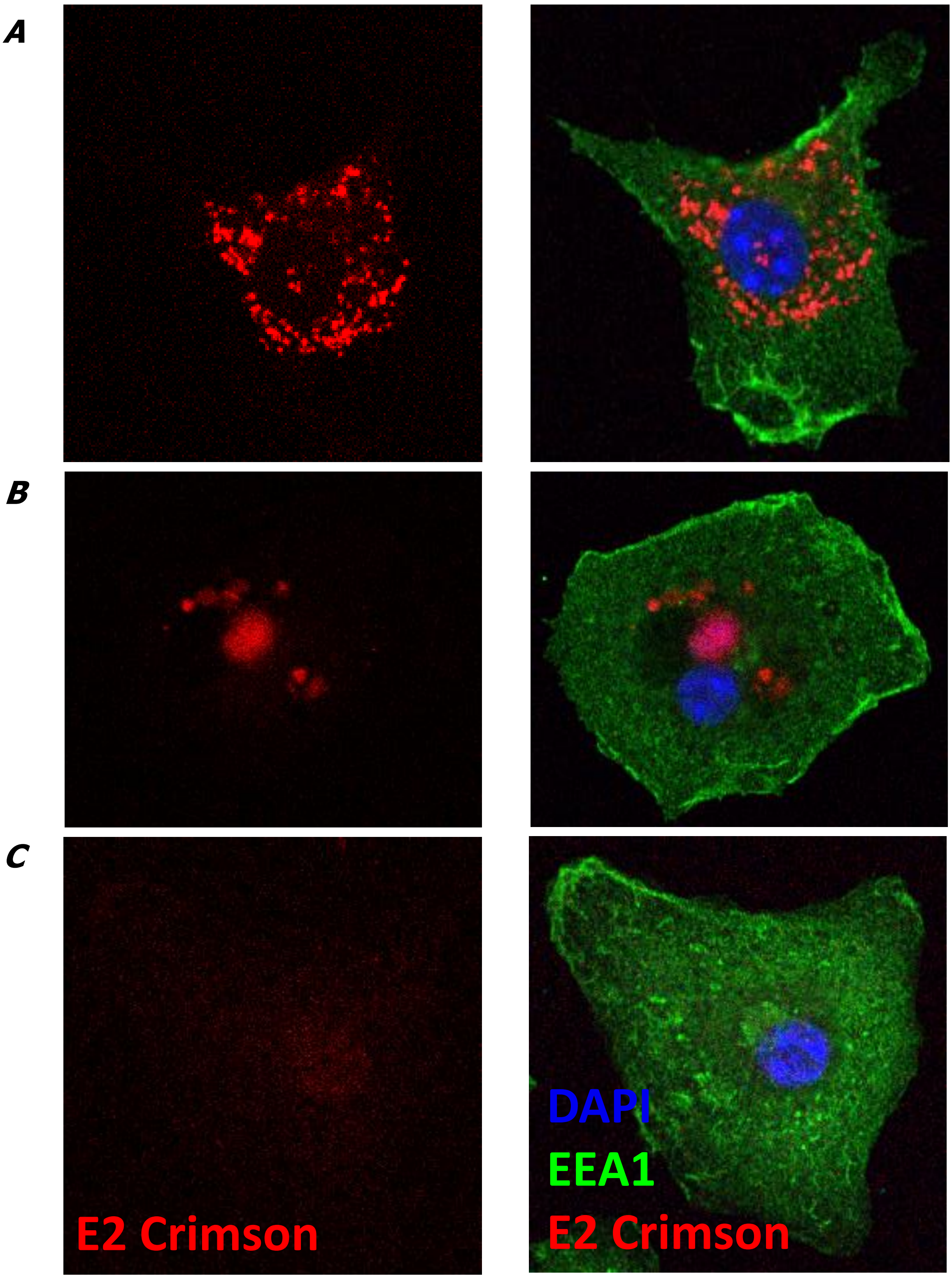


***Supplemental Figure 19.*** **Impact of Founder strain Background at Chr 10 and ADCP.** Bone Marrow Derived Macrophage CCxBALB/c (BMDM) in coculture with TUBO Crimson for 4 hours. E2-Crimson (red) is shown on the left and the 3-color image on the right (DAPI-blue, EEA1-green). Representative image of a BMDM differentiated from ***(A)*** CC035CF1, a predicted responder. ***(B)*** CC031CF1, a predicted responder. ***(C)*** CC024CF1, a predicted non-responder.

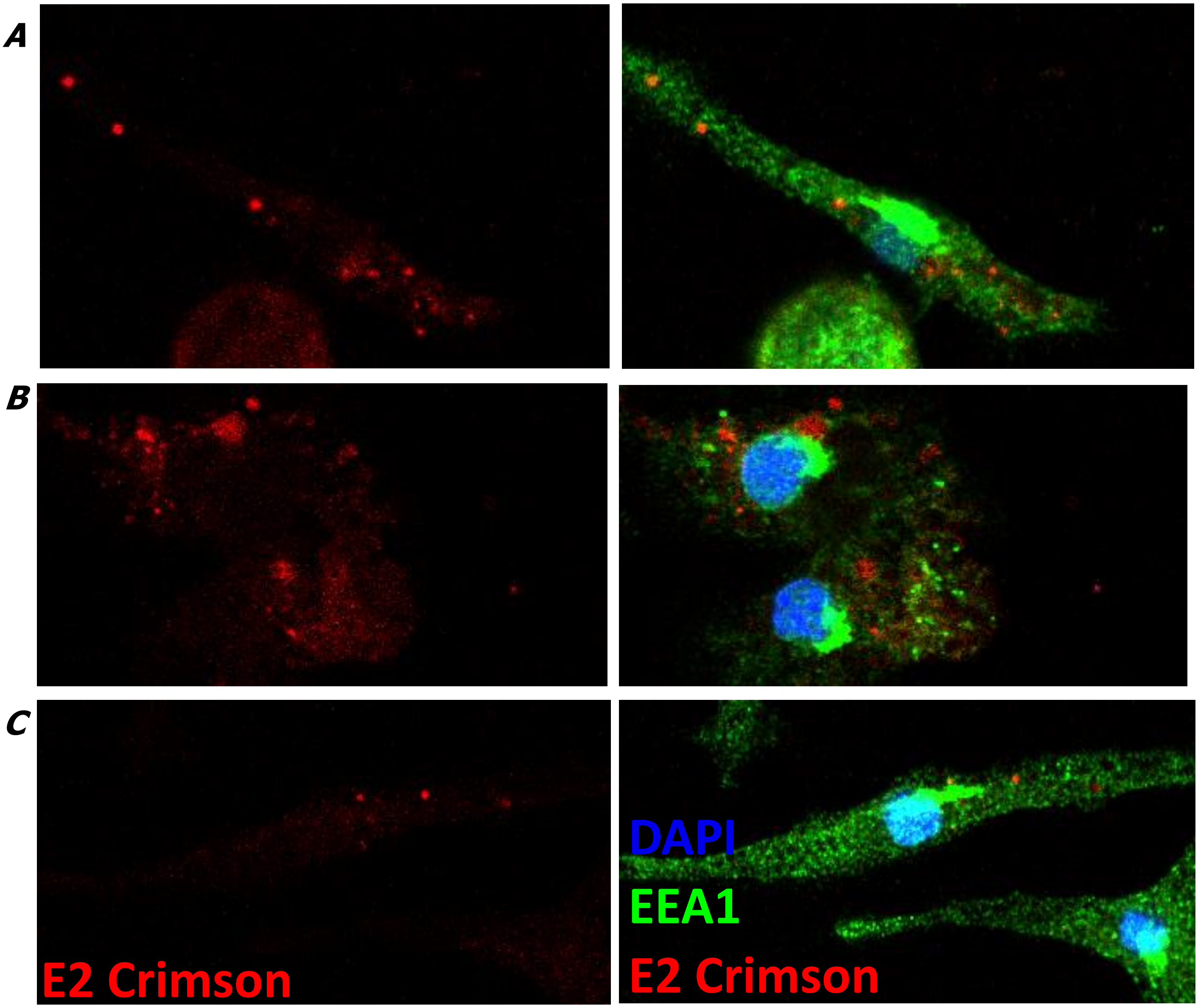


***Supplemental Figure 20.*** **Impact of Founder strain Background at Chr 10 and ADCP.** Bone Marrow Derived Macrophage CCxBALB/c (BMDM) in coculture with B16/Crimson for 4 hours. E2-Crimson (red) is shown on the left and the 3-color image on the right (DAPI-blue, EEA1-green). Representative image of a BMDM differentiated from ***(A)*** CC035CF1, a predicted responder. ***(B)*** CC031CF1, a predicted responder. ***(C)*** CC024CF1, a predicted non-responder.

A

B

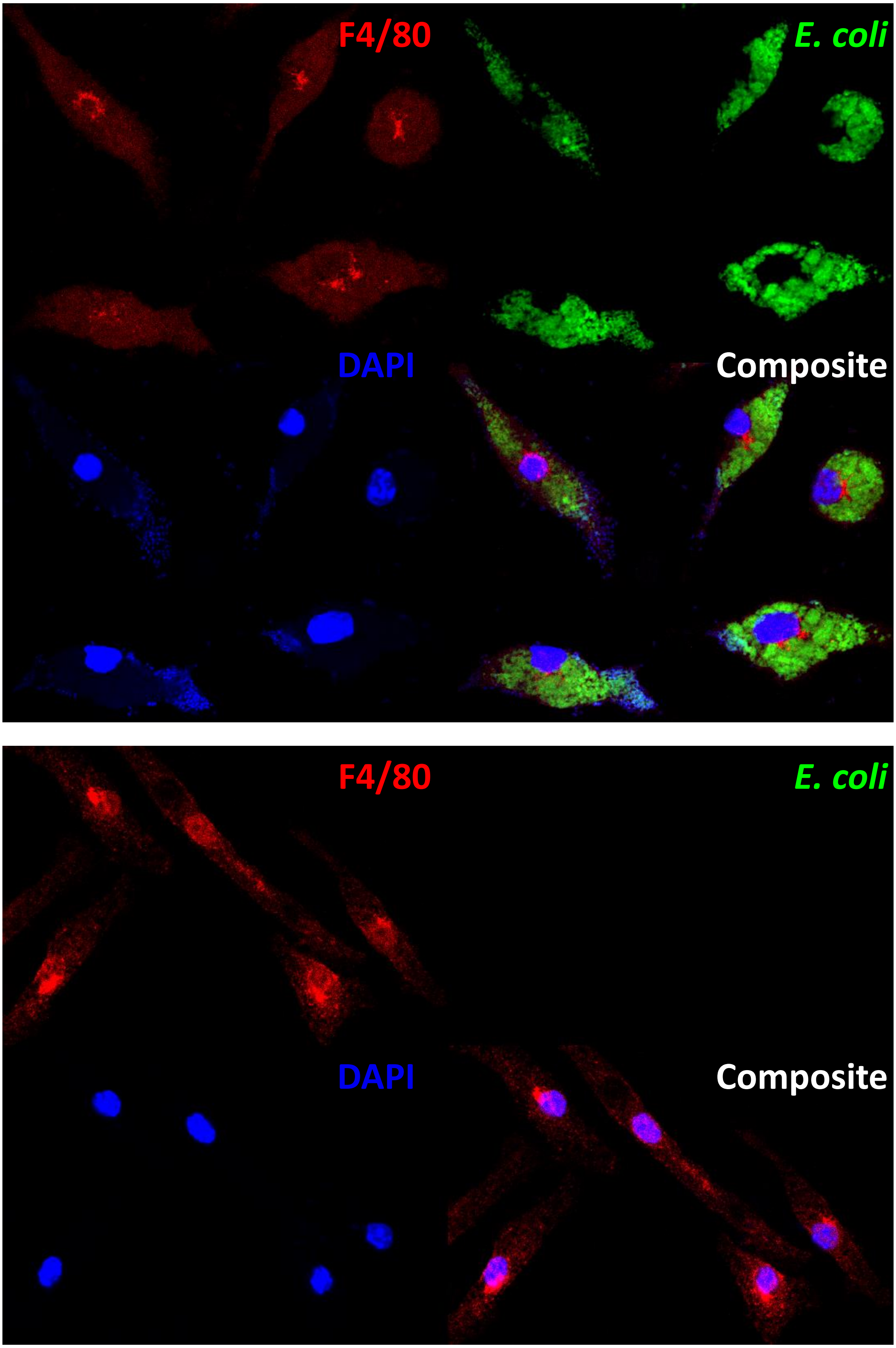


***Supplemental Figure 21.*** **Ab-independent phagocytosis is unaffected by genetic background at Chr 10.** Bone Marrow Derived Macrophage CC024CF1 in coculture with Fluorescein stained *E. coli*. Representative image of a BMDM differentiated from non-responder model. *(**A**)* CC024CF1 with *E.coli*. *(**B**)* CC024CF1, *E.coli* free control.

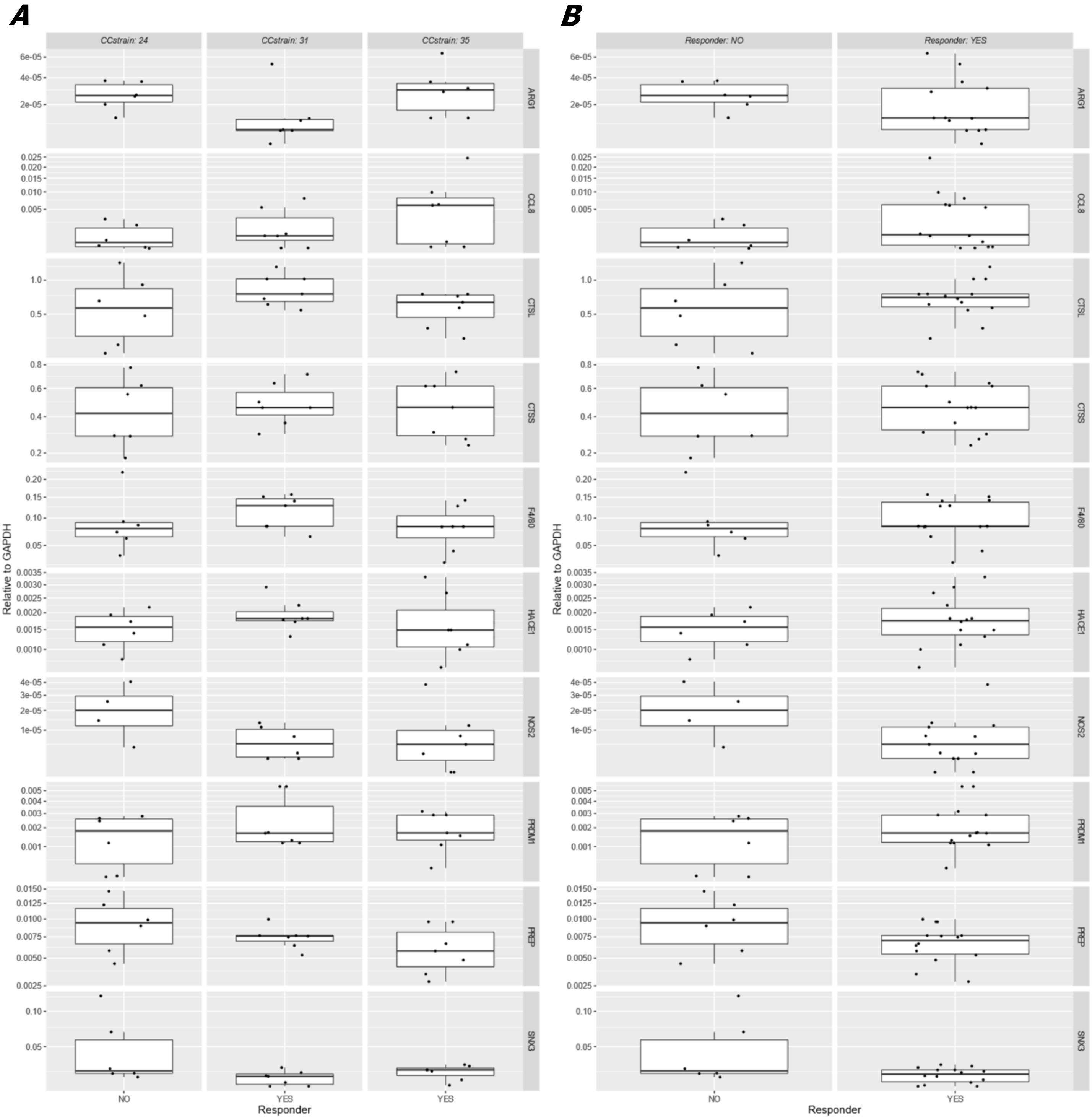


***Supplemental Figure 22.*** **qRT-PCR of various Chr 10 genes and canonical macrophage maturation and polarization genes.** BMDM in culture derived from CC024CF1, CC031CF1, and CC035CF1 mouse models split by (**A**) Model. (**B**) Response – CC024CF1 (No), CC031/35CF1 (Yes). All results shown are not significant by T-test.

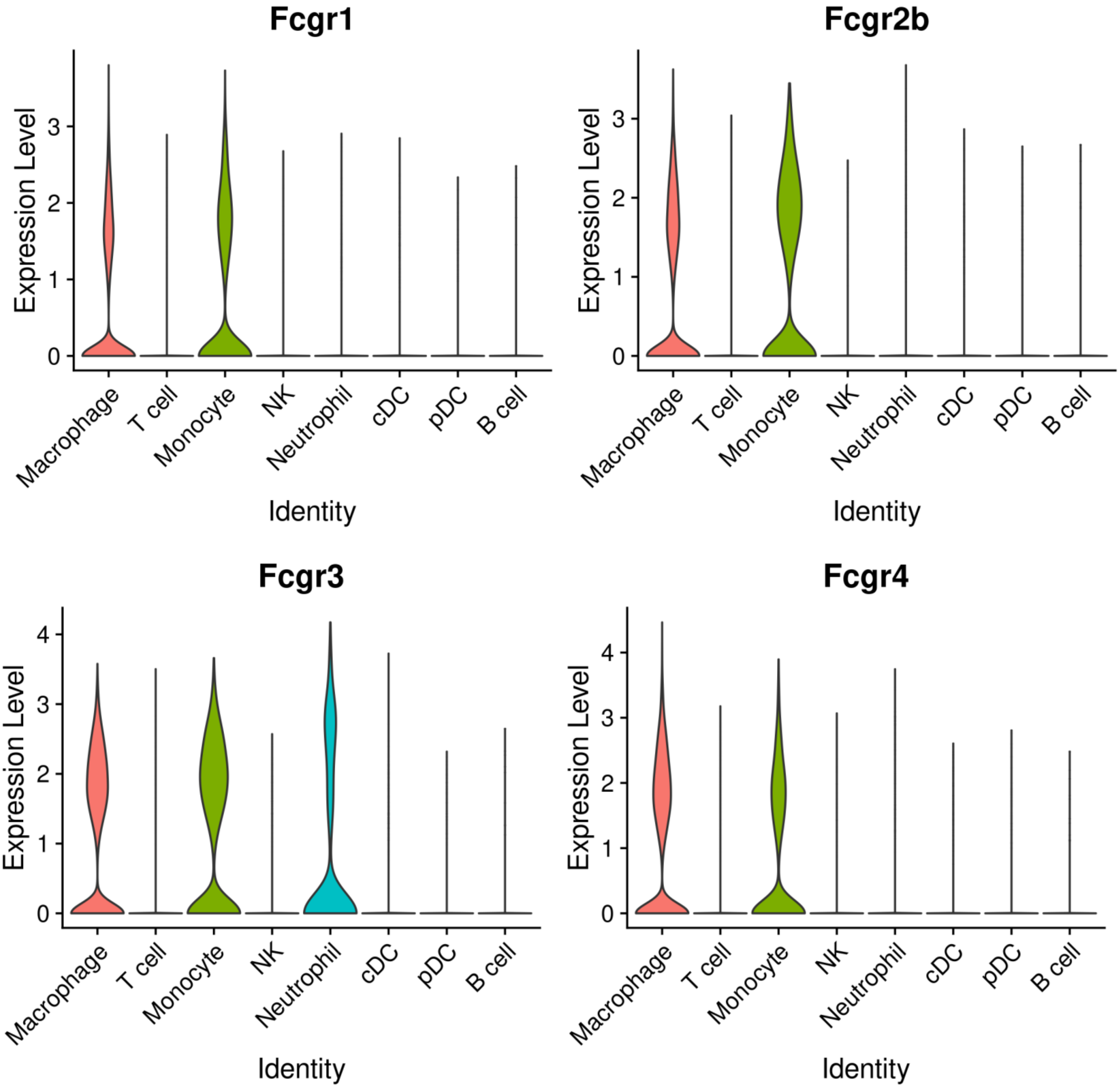


***Supplemental Figure 23.*** **$F_c$-receptor transcription implicates phagocytic cell types as the prime driver of targeted immunotherapy.** Expression of $F_c\gamma$-receptor across cell types via scRNA-seq.

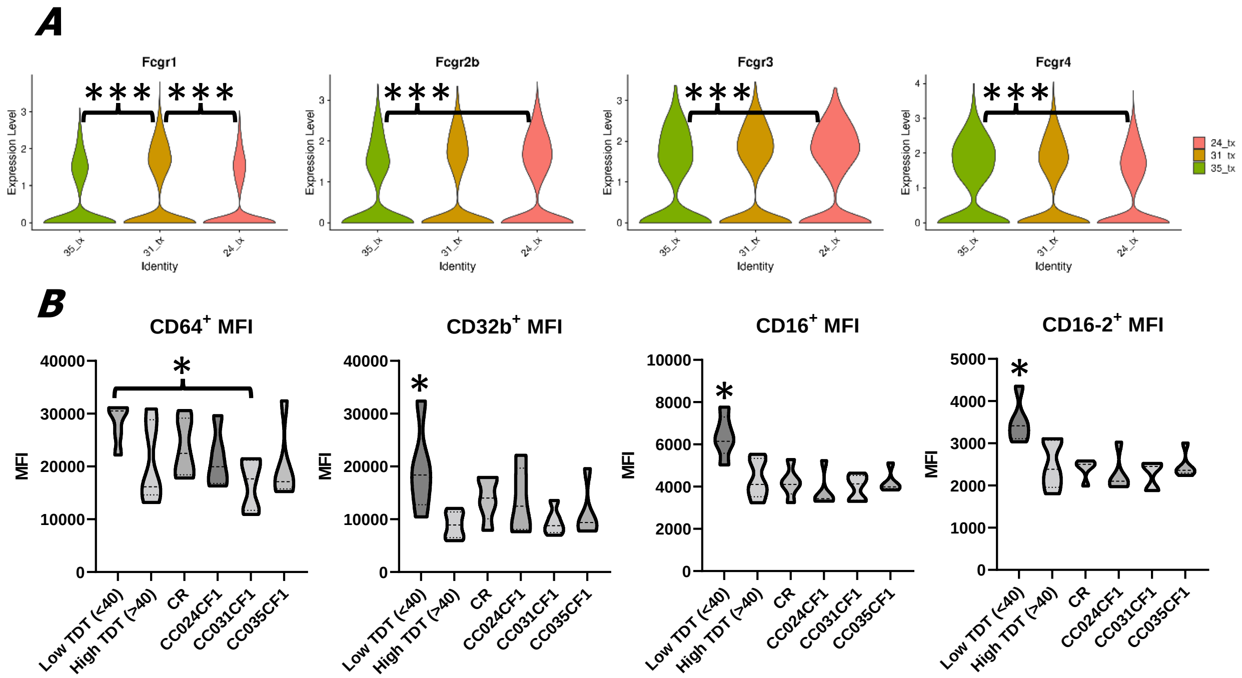


***Supplemental Figure 24*. Influence of genetic background on $F_{cR}$ diversity in the tumor microenvironment and on *ex vivo* BMDM.** *(A)* Tumor infiltrating Mφ expression of $F_cR$ transcripts by genetic background via scRNA-seq. (**B**) BMDM from DOCF1 and CCxCF1 mice stained for $F_cRs$ CD64, CD32, CD16, CD16-2. Mφ were gated on live, F4/80$^+$ cells.

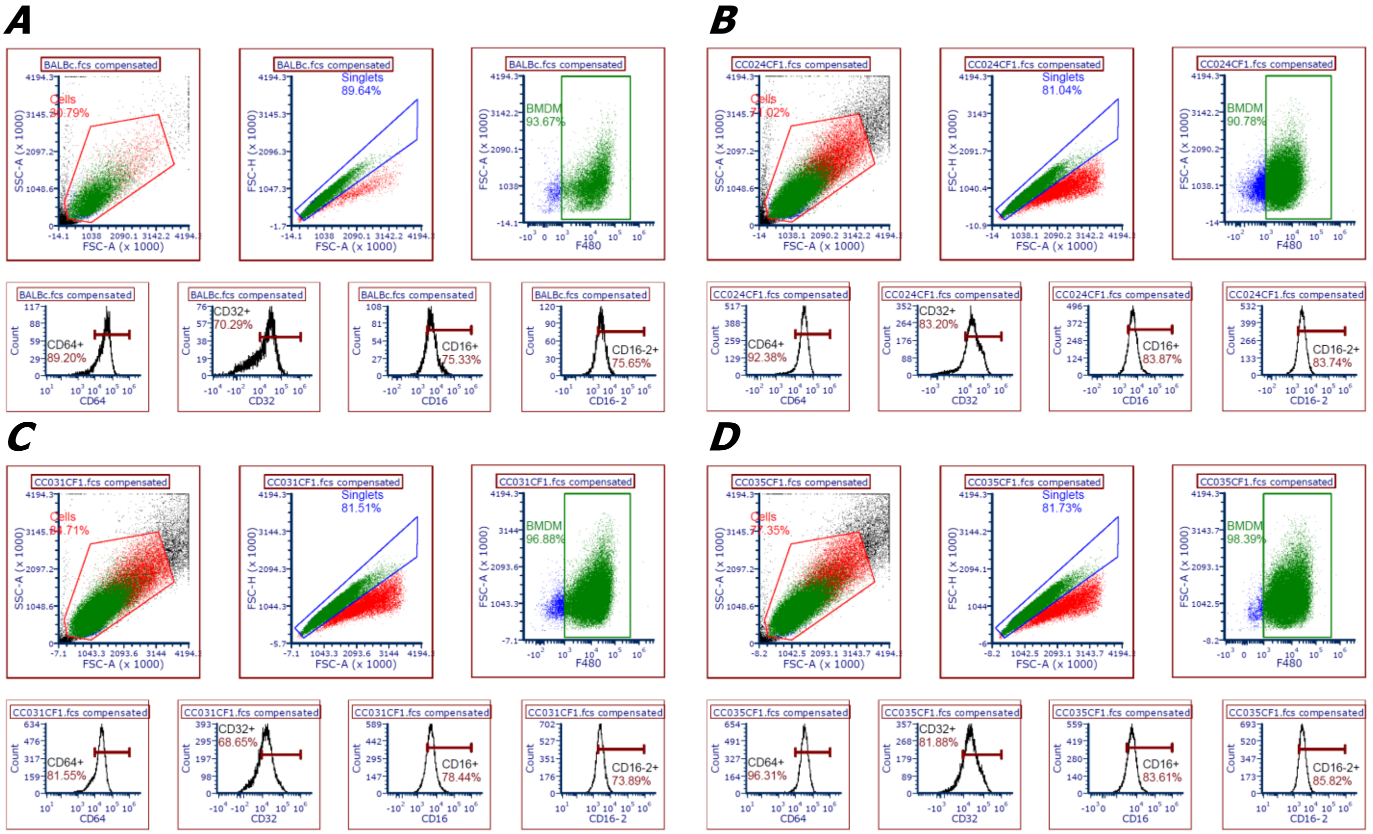


***Supplemental Figure 25*. Representative $F_c$ receptor diversity of BALB/c and CCxCF1 BMDM.**
***(A)*** BALB/c. ***(B)*** CC024CF1. ***(C)*** CC031CF1. ***(D)*** CC035CF1.
